# Scalable probe-based single-cell transcriptional profiling for virtual cell perturbation mapping and synthetic biology phenotyping

**DOI:** 10.64898/2026.02.04.703058

**Authors:** Jason T. Swinderman, Po-Yuan Tung, Aidan Winters, Laine Goudy, Caroline M. Wilson, Lexi R. Bounds, Noam Teyssier, Ayush Agrawal, Alex Dobin, Tony Hua, Hani Goodarzi, Felix Y. Feng, Alex Marson, Dave P. Burke, Patrick D. Hsu, Yusuf H. Roohani, Silvana Konermann, Michael Kosicki, Nianzhen Li, Luke A. Gilbert

## Abstract

Large-scale single-cell transcriptional phenotyping of genetic perturbations (perturb-seq) links genes to phenotypes and should enable virtual cell predictive modeling and cellular engineering. However, current perturb-seq single-cell methods are costly, information sparse and require barcodes for many applications. We developed ProPer-seq, a perturb-seq method that uses multiplexed custom DNA probe panels to measure and phenotype synthetic biology perturbations at single-cell resolution without barcodes, including multidomain proteins and sgRNAs. ProPer-seq faithfully reproduces gold-standard perturb-seq phenotypes while achieving 4-fold cost reduction and 50% increased gene detection per cell. As a scalable fixed-cell profiling method, ProPer-seq enables atlas-scale profiling for virtual-cell initiatives and demonstrates data quality suitable for training and validating predictive models. Lastly, ProPer-seq’s targeted detection of modular transgenes enables library-on-library perturbation profiling of combinatorial synthetic protein design spaces. We applied this to 3,550 sgRNA × dCas9 effector combinations as well as 260 CAR × ORF combinations dynamically profiled in primary T cells, revealing principles of transcriptional control and cell state modulation by multidomain synthetic transgenes.

## INTRODUCTION

Biological systems respond to genetic and environmental perturbations in a complex, cell- and context-specific manner, and exhibit redundancy that makes it challenging to discover underlying regulatory circuits and to achieve desired therapeutic outcomes^1–3^. Single cell functional genomics platforms, such as perturb-seq, provide a method to address these issues by enabling rich phenotypic readouts of perturbations at the single-cell level^4,5^. These approaches have so far enabled the study of such complex topics as the unfolded protein response, chimeric antigen receptor (CAR) T cell behavior following multiplex genetic perturbations and the impact of combinatorial enhancer perturbation^6–9^. Large scale perturb-seq training data are also fundamental to current efforts to create “virtual cell” models^1–3^.

Despite their potential, available perturb-seq approaches face significant limitations that restrict their scale, adoption and power to discover new biology^10^. Currently, the most widely used single cell RNA-sequencing approaches are unbiased, which results in a large fraction of sequencing capacity being deployed on lower information high-abundance transcripts^11–13^. This drives up the costs, especially for experiments in which lowly abundant, but highly informative transcripts, such as those coding for transcription factors, are of interest. Further, most single-cell RNA sequencing methods use a reverse transcriptase-based approach and capture only the 5’ or 3’ ends of the transcript, making it highly challenging to disambiguate natural or synthetic transcripts that share terminal sequences, for example alternatively spliced transcripts or CARs engineered with different combinations of similar immunomodulatory motifs. While this issue can be addressed by barcoding synthetic genes, it makes screening workflows more complex and unreliable. Using 5’ or 3’ readouts also significantly reduces the quality of data obtained from cells fixed with paraformaldehyde, which fragments the RNA, limiting the flexibility of screening approaches. Alternatively, probe-based methods for single cell transcriptome analysis that leverage custom DNA probe pools are emerging^14,15^. In theory, probe-based approaches should enable detection of internal RNA sequences in fixed cells; however, the sensitivity of these developing assays to accurately phenotype perturbations has not been established.

To address these limitations, here we develop and benchmark ProPer-seq, a **<u>pro</u>**be-based **<u>per</u>**turb-seq method combining probe-based, fixed-cell single-cell RNA sequencing with custom-designed probes to detect transgenes and/or single-guide RNAs (sgRNAs), enabling us to link perturbations to their transcriptional phenotype without barcodes. In a head-to-head comparison to standard 5’ and 3’ perturb-seq profiling methods, ProPer-seq detects more genes per cell at a substantially reduced cost, without sacrificing readout accuracy. Crucially, we demonstrate its ability to disambiguate synthetic transcripts at single cell resolution regardless of their 5’ or 3’ similarity, enabling the measurement of previously challenging synthetic perturbations without barcoding in genetically unlinked library-on-library perturbation experiments. To demonstrate the utility of ProPer-seq, we apply our method to diverse workflows, from profiling genetic interactions and perturbing gene regulatory elements with multi-domain dCas9-based effectors at low- or high-multiplicity of infection, to characterizing therapeutic transgene libraries in primary human T cells. Our results establish ProPer-seq as a scalable platform for virtual cell models^16^ and for systematic exploration of complex cellular perturbation phenotypes.

## RESULTS

### A ProPer-seq platform for large-scale single-cell perturbation profiling initiatives

The central challenge of perturb-seq assays is linking the identity of a genetic perturbation associated with gene knockout, knockdown or overexpression to a transcriptional phenotype at single-cell level. State-of-art perturb-seq approaches utilize 5’ or 3’-end RNA-sequencing methods based on reverse transcription, which do not natively capture non-polyadenylated RNA polymerase III-generated sgRNA transcripts identifying the perturbation^4,5^. Currently, this necessitates either direct capture of the sgRNA transcripts, incorporation of the sgRNA into a polyadenylated RNA polymerase II-generated transcript (CROP-seq) or linking a transcribed barcode with each sgRNA^7,17,18^. These approaches suffer issues such as loss of linkage between barcode and sgRNA due to lentiviral template switching and increase experimental complexity. Probe-based single-cell RNA-sequencing in theory would enable low-cost, high transcriptome resolution, custom RNA-profiling workflows, but has not yet been benchmarked in perturb-seq contexts^15^. We speculated that a probe-based approach would enable a unified perturb-seq framework that natively captures sgRNAs, while reducing the overall cost, increasing gene detection and enabling scalable complex synthetic biology experiments.

We designed an experiment to benchmark state-of-art 5’ and 3’ direct-capture perturb-seq workflows to a probe-based perturb-seq platform (**Figure 1A**). Based on our prior experience using perturb-seq assays, we selected sgRNAs designed for CRISPR activation (CRISPRa) targeting promoters of 42 transcription factor genes along with negative control sgRNAs (sgNeg) and cloned matched lentiviral sgRNA libraries with and without sgRNA capture sequences. We infected each sgRNA library at low multiplicity of infection (MOI) into a population of K562 cells that stably express a CRISPRa transgene, sorted infected cells to purity and profiled at single-cell resolution the transcriptional phenotypes induced by gene activation 6 days post-transduction. Specifically, we compared cells infected with sgRNAs lacking capture sequences, profiled using 5’ direct-capture RNA-seq or probe-based RNA-seq, and cells infected with sgRNAs with capture sequences, profiled using 3’ direct-capture RNA-seq (**Figure 1A**). To generate a robustly sampled dataset to compare performance we profiled each method at high cell and read coverage (357-743 cells per CRISPRa perturbation, 10,000-13,500 reads per cell on the average; Methods).

**Figure 1.**
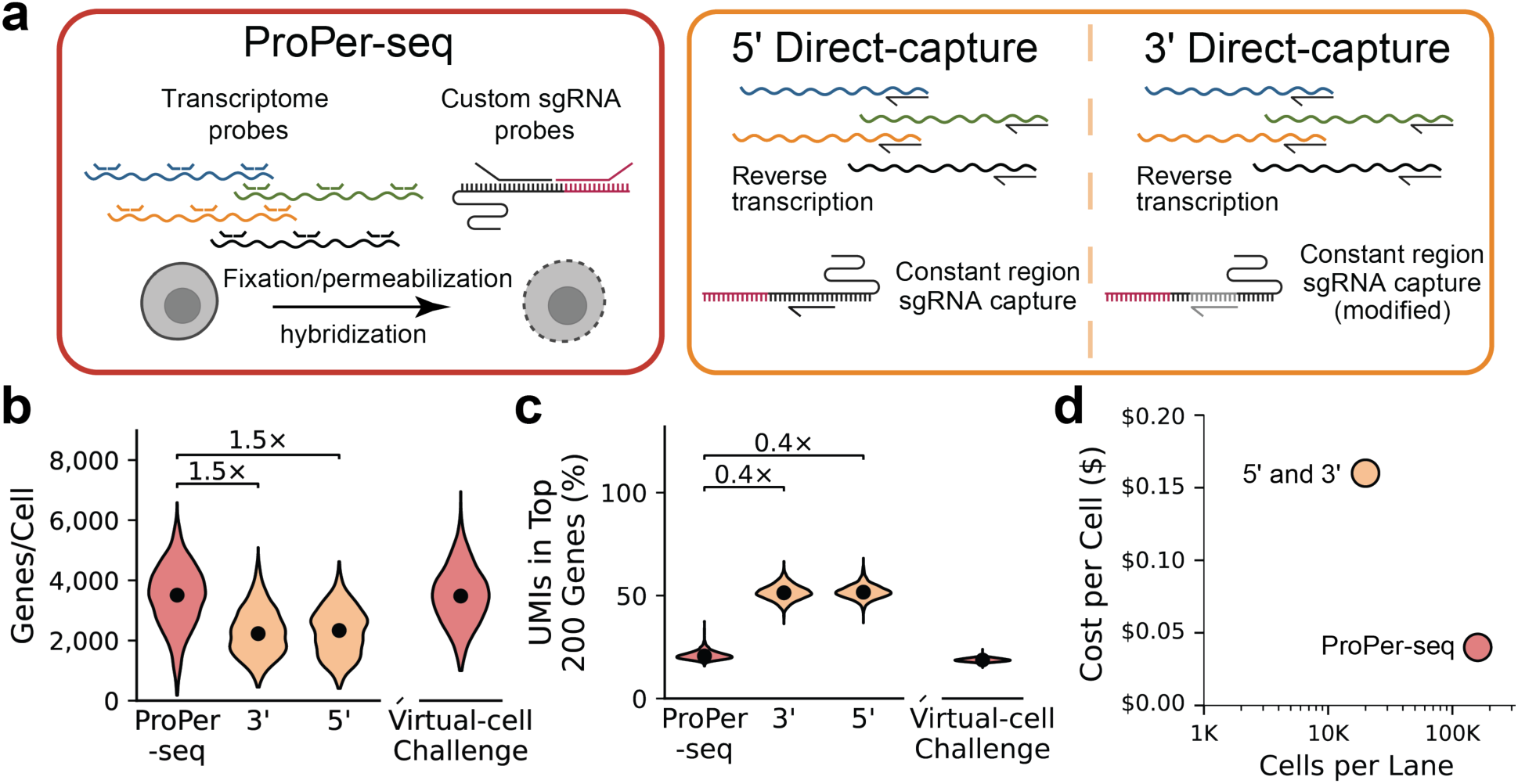
Scalable probe-based perturb-seq with ProPer-seq. **a.** Overview of benchmarked techniques. Left: fixed-cell probe-based single-cell RNA-seq per-turb-seq (ProPer-seq) with custom sgRNA detection, right: conventional 5’ perturb-seq and 3’ perturb-seq protocols using un-fixed cells and reverse transcription for generation of single-cell transcriptome and sgRNA libraries. **b.** Violin plots of the number of genes detected per cell in K562 cells (ProPer-seq, 3’, 5’) or human embryonic stem cells from Arc Virtual-Cell Challenge dataset^16^ assigned sgNeg (negative control sgRNA). Cell and counts were downsampled evenly for each method (N = 2,500 sgNeg cells). **c.** Violin plots of the fraction of UMIs in top 200 highest expressed genes. Same data as in (b). **d.** Cost of library construction per cell and the number of cells passing quality control per lane for ProPer-seq compared to 5’ perturb-seq and 3’ perturb-seq.

To measure the sgRNAs present in each cell using the probe-based approach, we designed a custom sgRNA probe library that was spiked into the transcriptome probe library (see Methods). Our probe design for sgRNA capture utilizes a custom 5’ probe that hybridizes to the sgRNA unique protospacer sequence and a common 3’ probe that binds to the sgRNA constant sequence which can be ligated, amplified and then counted by sequencing **(Figure 1A)**.

We examined the validity of ProPer-seq as compared to 5’ and 3’ perturb-seq experiments in terms of baseline measurement of gene expression, sgRNA capture rates and perturbation phenotypes. We found that despite ProPer-seq having fewer sgRNA UMIs on average per cell (ProPer-seq: 48 umi/cell, 3’: 175 umi/cell, 5’: 1,524 umi/cell), overall cell assignment rates and proportionate sampling of each sgRNA was comparable across methods (**Extended Data Fig. 1A;** ProPer-seq vs 5’-direct capture χ^2^=90.3, p = <0.01, r = 0.99). Additionally, we found the degree of target gene activation in sgRNA-assigned cells compared to sgNeg to be consistently measured in ProPer-seq and direct-capture approaches (**Extended Data Fig. 1B**). To understand whether gene sampling between fixed probe-based assays recapitulated live cells processed with reverse-complement based RNA detection we sampled cells expressing negative control sgRNAs (sgNeg) at an equal read depth and found largely consistent distribution of reads (**Extended Data Fig. 1C**). To then compare whether ProPer-seq was detecting similar transcriptional content as 5’ and 3’ direct-capture we compared our cells processed by ProPer-seq, 5’, and 3’ as well as the recently released Virtual Cell Challenge training dataset that we generated with ProPer-seq in H1 embryonic stem cells^3,16^. In doing so we found that probe-based transcript detection resulted in increased genes measured per cell, while also sampling highly abundant transcripts less frequently thus providing richer data per cell which will benefit training virtual cell models (**Figure 1B-C**). These trends were even more pronounced when considering only coding-genes (**Extended Data Fig. 1D-F**). Lastly, ProPer-seq experiments can process far more cells per lane at lower cost per cell both of which enable large scale perturb-seq experiments required for virtual cell training efforts (**Figure 1D**).

Having established the feasibility of ProPer-seq, we designed a second larger sgRNA library to measure transcriptional phenotypes associated with activation of 83 genes previously described in a widely benchmarked 3’ perturb-seq dataset^19^. The same pool of perturbed cells was processed in parallel with ProPer-seq and 5’ perturb-seq workflows. We examined CRISPRa induced on-target gene activation and the associated perturb-seq phenotypes arising from transcription factor overexpression. The degree of target gene activation measured by 5’ perturb-seq and ProPer-seq was again highly similar showing our ability to measure target gene transcriptional perturbations is robust across approaches (**Figure 2A**). Next, we evaluated the transcriptional phenotypes induced by gene overexpression using energy-distance scores, a well-established single cell phenotyping metric that uses the difference between negative control transcriptomes and the gene perturbation transcriptomes to score phenotypes^10^. We observed a strong correlation between energy-distances derived from each perturb-seq method across perturbations (**Figure 2B**). To comprehensively compare how ProPer-seq and 5’ perturb-seq corresponded and to evaluate our ability to recapitulate our previously published results, we plotted the perturbation phenotypes of all gene activation perturbations comparing our ProPer-seq and 5’ perturb-seq data to published 3’ perturb-seq data^19^. We found highly replicable phenotypes that were comparably measured by ProPer-seq and 5’ perturb-seq approaches relative to the published 3’ perturb-seq data (**Figure 2C-D**). This demonstrates that perturb-seq is robust across methods, individuals and time which should be enabling for large scale projects. We conclude that ProPer-seq is comparable to current state-of-art 5’ and 3’ perturb-seq methods but has several key advantages due to information content per cell, use of fixed cells, scalability and cost.

**Figure 2.**
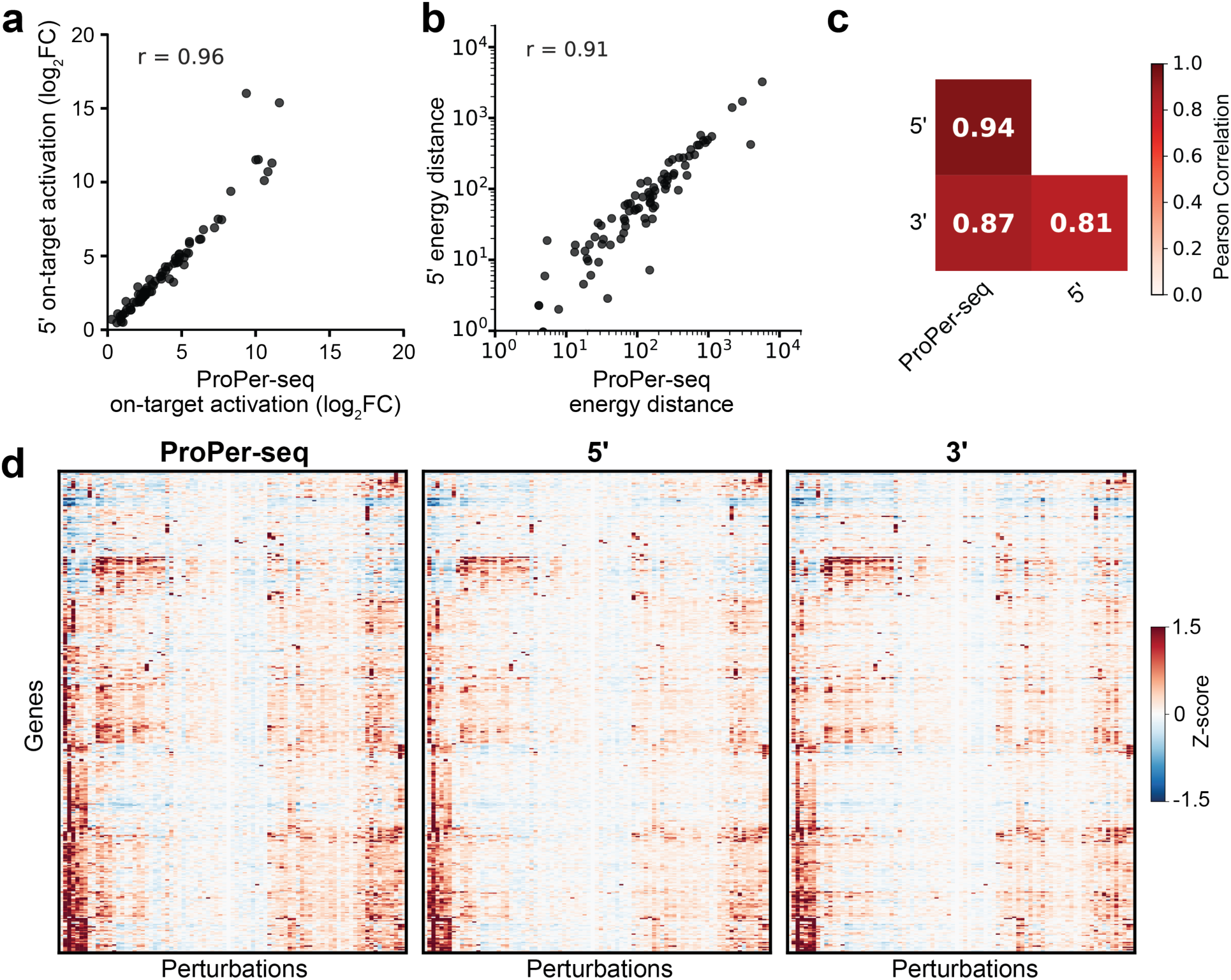
Perturbation fidelity with ProPer-seq. **a.** On-target gene activation by 84 CRISPRa perturbations measured with 5’ perturb-seq or ProPer-seq. Activation is expressed as log_2_ fold-change of target gene expression in cells assigned a cognate sgRNA compared to cells assigned sgNeg (Pearson’s r = 0.96, p-value < 0.01, N = 84 pseudobulk single sgRNA perturbations). **b.** Perturbation effect size calculated as energy-distance^10^ from 1,000 permutations of CRISPRa perturbations compared to cells assigned sgNeg measured with 5’ perturb-seq or ProPer-seq (Pearson’s r = 0.91, p-value < 0.01, N = 84 pseudobulk single sgRNA perturbations). **c.** Pearson correlations between transcriptional effects of 84 CRISPRa perturbations on 4,088 consensus overlapping highly variable genes measured with 5’ perturb-seq (N= 28,268 cells), 3’ perturb-seq (N = 54,176 cells) or ProPer-seq (N = 21,187 cells). **d.** Transcriptional effects of 84 CRISPRa perturbations on 4,088 consensus overlapping highly variable genes measured with 5’ perturb-seq, 3’ perturb-seq or ProPer-seq.

### ProPer-seq enables robust quantification of multigene perturbations

Having demonstrated the robustness of ProPer-seq in single-gene perturbation experiments, we applied this workflow to genetic-interaction experiments requiring multiplexed sgRNA detection at the single cell level. We have previously shown that transcriptomic quantification of genetic-interactions (GIs) between pairs of perturbed genes requires accurate and robust assignment of combinations of sgRNAs at single-cell resolution as well as rich transcriptomic measurements^19^. We set out to test whether ProPer-seq sgRNA detection and transcriptional phenotyping was sufficiently robust to enable measurements of genetic interactions induced by CRISPRa activation of gene pairs. To investigate, we designed a dual-sgRNA library containing 193 unique dual-sgRNA constructs, reflecting 109 dual-gene GI pairs and 83 single-gene targeting constructs **(Figure 3A)**. Out of 109 GIs, 27 were previously measured GIs^19^. To select the remaining novel gene targeting pairs, we had previously trained GEARS^20^, a predictive deep learning approach, on select single gene perturbation perturb-seq data^19^, and predicted all possible pairwise GIs. However, none of these predicted GIs were experimentally tested^20^ leaving it unclear if perturb-seq data quality is robust enough to train deep learning models that accurately predict GIs which can be subsequently validated. We prioritized 82 previously unmeasured GIs across a range of predicted interaction phenotypes and primary fitness effects (**Extended Data Fig. 2A-B, Extended Data Table 1**). We balanced the sgRNA library according to measured fitness scores and infected the dual-sgRNA GI library at low MOI into a population of K562 cells that stably expressed a CRISPRa transgene, sorted infected cells to purity and profiled the transcriptional phenotypes by 5’ direct-capture or ProPer-seq 6 days after infection (**Figure 3A, Extended Data Fig. 2C**).

**Figure 3.**
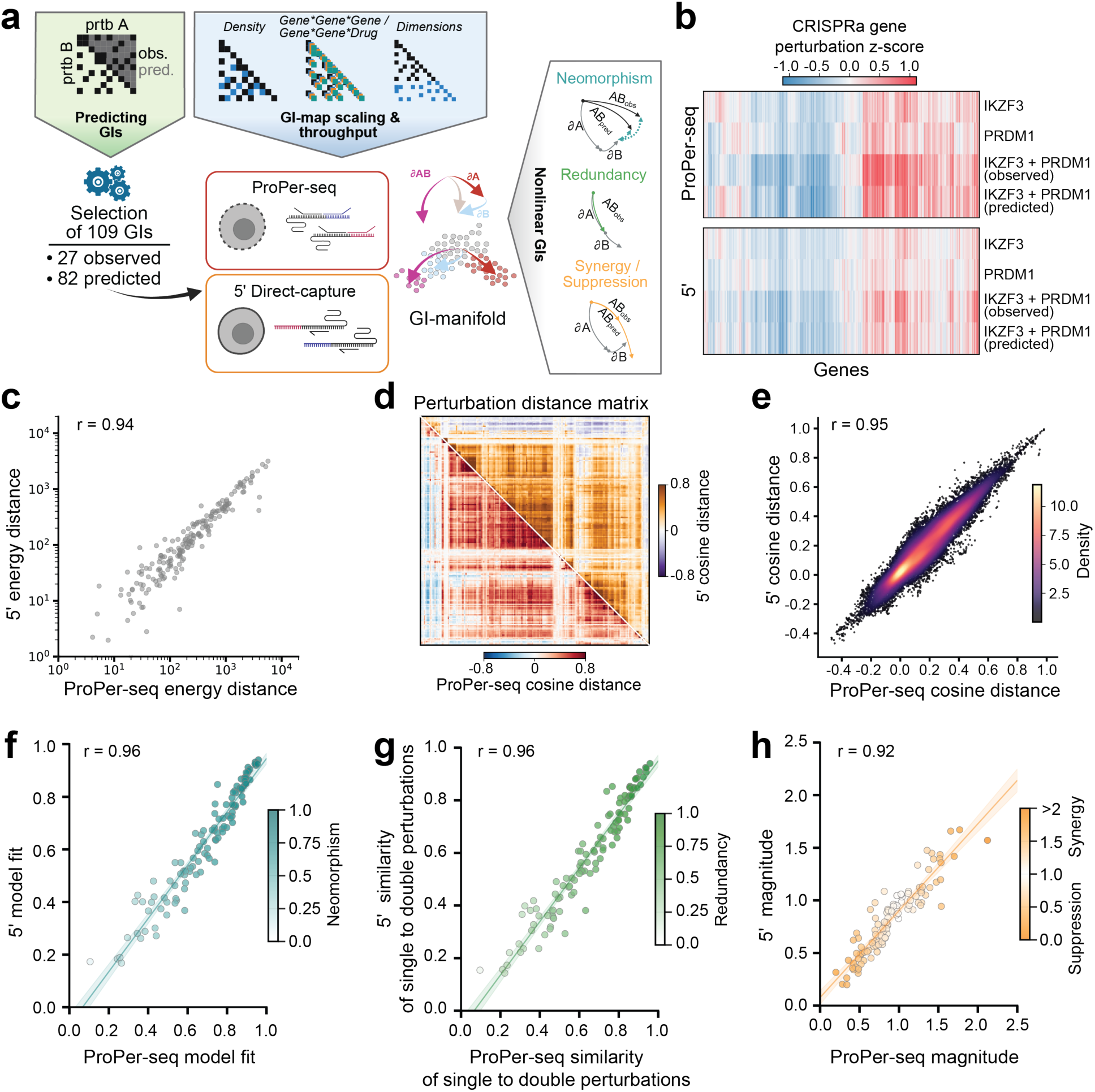
ProPer-seq enables robust quantification of multigene perturbations. **a.** Design of the genetic interaction (GI) experiment. Previously measured sparse genetic-inter-action perturbations^19^ in K562 CRISPRa were used to train a complete pairwise matrix of GIs with GEARS^20^. From these, 109 GIs were selected for experimental validation using ProPer-seq and 5’ perturb-seq, including 27 previously observed and 82 predicted GIs. Measured GIs were fit to a linear interaction model^19^ (see methods) to calculate distinct classes of nonlinear interac-tions including neomorphic, redundant, and synergistic GIs. **b.** An example of the observed and linear model predicted transcriptional impacts of combinato-rial CRISPRa perturbations. The impact of sgRNAs targeting IKZF3 and/or PRDM1 on consen-sus highly variable genes is shown, as measured by ProPer-seq (top) and 5’ perturb-seq (bot-tom). **c.** Perturbation effect size calculated as energy-distance^10^ from 1,000 permutations of CRISPRa perturbations compared to cells assigned sgNeg measured with 5’ perturb-seq or ProPer-seq (Pearson’s r = 0.94, p-value <0.01, N = 109 unique pseudobulk perturbations). **d.** Similarity of all measured perturbation effects shown by cosine-distance matrix of perturba-tion z-score. 5’ perturb-seq clustering (top) was applied to order ProPer-seq perturbations (bot-tom) **e.** Density scatterplot of cosine-distance values for all perturbation-perturbation pairs comparing ProPer-seq and 5’ perturb-seq (Pearson’s r = 0.95 p-value <0.01). **f-h.** Correlation plots of primary GI fit parameters; neomorphism (**f**), redundancy (**g)**, and syn-ergy/suppression (**h)** derived from model-fit, similarity of single to double perturbations, and magnitude of GI linear model as previously described^19^. Shown with fit curve and 95% confi-dence interval (Pearson’s r = 0.96, 0.96, and 0.92 for **f-h** respectively).

We observed robust detection of expected pairs of sgRNAs encoded by our sgRNA library in single cells by ProPer-seq with a similar number of useable cells relative to 5’ perturb-seq (**Extended Data Fig. 2C**). We also observed comparable on-target gene activation for each perturbation profiled by ProPer-seq and 5’ perturb-seq (**Extended Data Fig. 2D**). When evaluating perturbation effects against unperturbed control cells, both permuted energy-distance metrics and differential gene expression quantification, revealed high inter-assay consistency (r = 0.94 and r = 0.97, respectively, **Figure 3C, Extended Data Fig. 2E**). Next, we measured and modeled comparable interaction profiles for a representative additive GI (**Figure 3B**) and synergistic GI (**Extended Data Fig. 2F**). Consistent with our published 3’ perturbation phenotypes^19^, both single and dual gene CRISPRa perturbations were sufficient to produce strong changes in cell state. We extend this in our current data to show induction of erythroblast associated genes preferentially in GEARS predicted erythroblast GIs (**Extended Data Fig. 2G**) and produce consistent differentiation states including myeloid- and erythroid- associated transcriptional profiles (**Extended Data Fig. 2H-I**). To elucidate the distribution of perturbation-induced transcriptional responses, we implemented hierarchical clustering on a cosine distance correlation matrix and compared the concordance of pairwise cosine similarity across all perturbations. This analysis revealed similar correlation across all perturbation pairwise relationships as phenotyped by ProPer-seq and 5’ perturb-seq (r = 0.95, **Figure 3D-E**).

To determine whether ProPer-seq could quantitatively measure genetic interactions, we applied a linear model of single perturbations to quantify nonadditive genetic interactions as has been previously described (**Figure 3A**)^19^. Using model fit parameters, we compared the stratification of genetic interactions across four primary nonadditive classes: neomorphic, redundant, synergistic, and suppressive combinations. Our analysis captured a diverse set of GIs with model fit term distributions consistent with established findings. ProPer-seq and 5’ perturb-seq GI model fits similarly recapitulated originally calculated 3’ perturb-seq GI model parameters: model fit (Pearson’s r; 0.62 5’ perturb-seq, 0.66 ProPer-seq), magnitude (Pearson’s r; 0.66 5’ perturb-seq, 0.68 ProPer-seq), and similarity of single perturbations to dual (Pearson’s r; 0.61 5’ perturb-seq, 0.66 ProPer-seq. **Extended Data Fig. 3A-C**). Notably, all terms quantifying the primary genetic interaction classes showed high consistency across gene-pairs between ProPer-seq and 5’ perturb-seq (correlation coefficients for model fit, similarity of single to dual gene perturbation, and magnitude were 0.96, 0.96, and 0.92, respectively, **Figure 3F-H**). The robust concordance of global perturbation magnitudes and specific interaction fit terms across modalities validates the reproducibility of these approaches for measuring combinatorial perturbations and their extension to additional modalities enabled by probe-based transcript detection. Our classification of GIs with GEARS achieved strong performance metrics in categorizing GIs, with precision values of 0.87 for neomorphic, 0.93 for suppressive, and 0.65 for redundant interactions (**Extended Data Fig. 3D-J**). Together, our results demonstrate ProPer-seq workflows provide a scalable and efficient system for measuring genetic interactions, which—particularly when combined with machine learning prediction of interactors—offers a powerful solution for phenotyping genetic interactions, an innately exponential biological challenge.

### ProPer-seq enables genetically unlinked library-on-library CoMPoSE experiments

Having demonstrated the robustness and scalability of ProPer-seq for pooled multigene perturbation studies, we hypothesized that custom probe design could similarly enable pooled perturbation studies using synthetically engineered proteins. To this end, we designed a <u>Co</u>mbinatorial <u>M</u>odular <u>P</u>erturb-Seq <u>o</u>f <u>S</u>ynthetic <u>E</u>ffectors (CoMPoSE) experiment **(Figure 4A)** in which we perturbed cells with combinations of dCas9-based artificial transcription factors (“dCas9 effectors”) and sgRNAs targeting gene promoters. We refer to experiments that use two or more such unlinked and unbarcoded perturbations per cell as library-on-library experiments.

**Figure 4:**
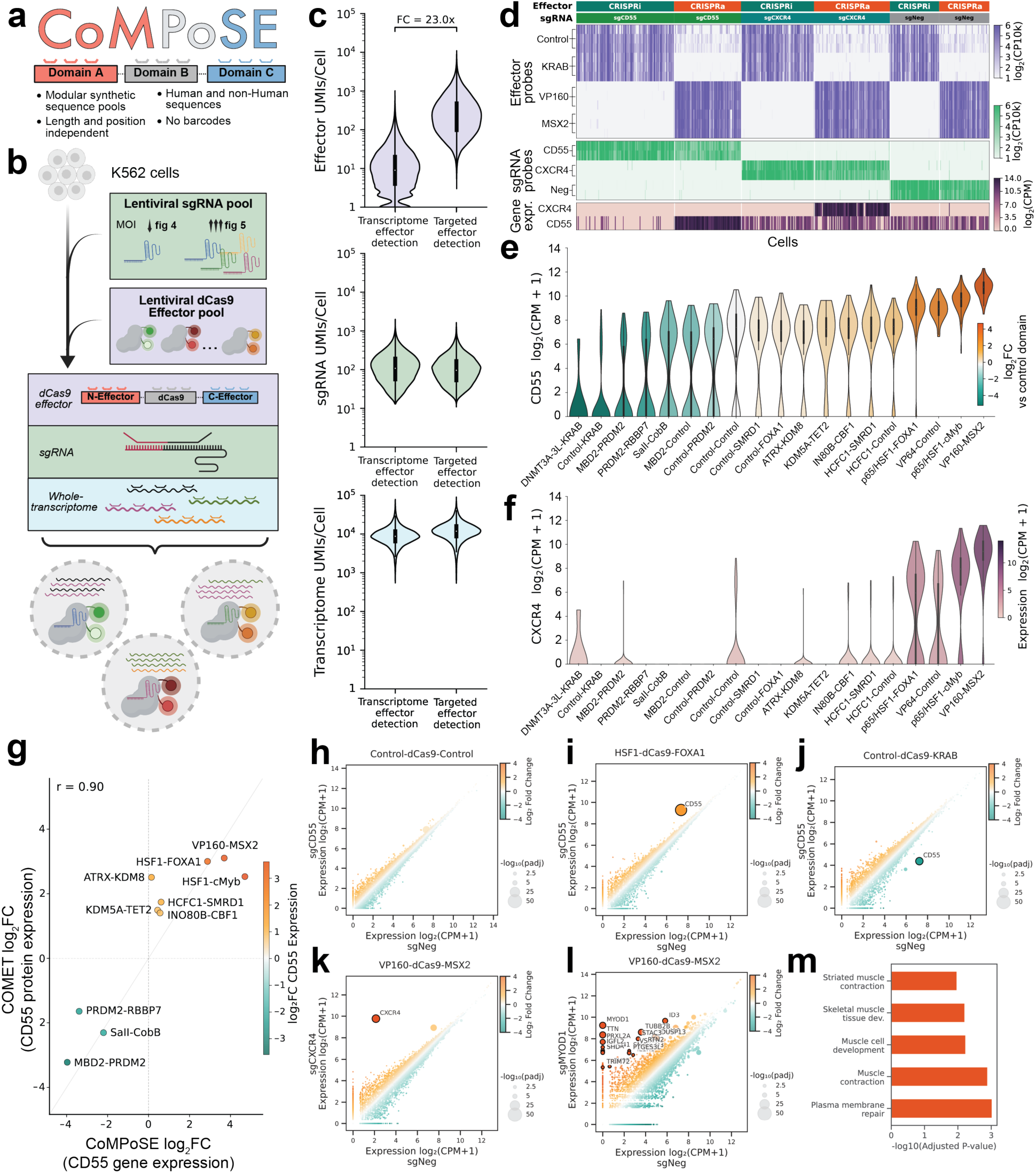
ProPer-seq enables genetically unlinked library-on-library CoMPoSE experi-ments. **a.** Design for Combinatorial and Modular Perturb-seq of Synthetic Effectors (CoMPoSE). Modu-larly designed transgenes are detected in single cells by probes tiling throughout the synthetic transgene transcript. **b.** Experimental schematic of CoMPoSE library-on-library experiments with sgRNA and dCas9 effector perturbations. Pooled probe hybridization allows for simultaneous construction of custom sgRNA and dCas9 effector sequencing libraries along with whole-transcriptome probe amplification. **c.** UMIs per cell for effectors (top), sgRNA (middle), and transcriptome (bottom) with or without targeted amplification of dCas9 effector pools. **d.** Heatmaps of single-cell detection of effector and sgRNA custom probes, and target (*CD55*, and *CXCR5*) expression. Grouped by sgRNA (sgNeg, sgCD55, and sgCXCR4) and effector (CRISPRi: Control-dCas9-KRAB and CRISPRa: VP160-dCas9-MSX2) assignment. **e-f.** Violin plots of all dCas9 effectors ordered by CD55 log_2_(fold-change) over control effector, showing for cells assigned each effector and either CD55 (**e**) or CXCR4 (**f**) sgRNAs. Plotted as the log_2_CPM of the target gene on the y-axis and colored by the log_2_(fold-change) against a control dCas9 effector (**e**) or log_2_(CPM) (**f**). **g.** Scatterplot of dCas9 effector activity measured as log_2_(fold-change) enrichment of CD55 high or low effectors from COMET CD55 surface protein-based screen^21^ plotted against CoM-PoSE measured modulation of CD55 transcript detection (linear fit Pearson’s r = 0.80, p-value < 0.01). **h-j. S**catter plot of all detected gene expression in control effector (**h**), CRISPRa (**i**), or CRISPRi (**j**) containing cells with sgCD55 or sgNeg. Colored by log_2_ fold-change and with point-size pro-portional to pseudobulk DESeq2 calculated -log_10_(padj). **k,l.** Whole-transcriptome scatter plot of gene expression in cells with a CRISPRa effector com-paring sgNeg to sgCXCR4 (**k**), or sgMYOD1 (**l**). Colored by log_2_(fold-change) and with point-size proportional to pseudobulk DESeq2 calculated -log_10_(padj). **m.** Gene Ontology pathway enrichment for significantly upregulated genes — adjusted p-value < 0.05 and log2(fold-change) > 0.5 — with sgMYOD1.

Specifically, we cloned a lentiviral library of 9 dual-sgRNA constructs, where each construct encodes a pair of sgRNAs that targets the same promoter or encodes a pair of negative control sgRNAs, co-expressing BFP. We also cloned a second lentiviral library of 19 dCas9 effectors (co-expressing mNeonGreen) that we had previously validated as capable of repressing or activating transcription when targeted to the CD55 or CXCR4 gene (**Extended Data Fig. 4A**)^21^. Each dCas9 effector was designed with a dCas9 domain flanked by an N-terminal and a C-terminal transcriptional effector domains.

We transduced K562 cells with the sgRNA library at low MOI and then 2 days later with the effector protein library at low MOI such that across the population of infected cells we expected 171 combinations of library-on-library infected cells (**Figure 4B**). We used FACS to isolate cells positive for sgRNA construct integration and dCas9 effector expression (BFP+/mNeonGreen+), and proceeded to ProPer-seq after 5 days, to allow effector associated transcriptional profiles to manifest (**Extended Data Fig. 4B**). Specifically, paraformaldehyde-fixed cells were hybridized with a pool of three probe sets: custom dual-sgRNA-library probes, custom probes targeting dCas9-effector library N- and C-termini and the transcriptome probe library (**Figure 4B, Extended Data Fig. 4C-D**). We designed custom dCas9 effector probes to minimize nonspecific transcriptome binding and maximize effector probe efficiency. To increase effector signal, we selectively amplified effector probes and sequenced the resulting library (see Methods, **Extended Data Fig. 4E-G**). We observed a substantially enhanced effector signal without impact on sgRNA detection or gene expression compared to the nonselective library amplification (**Figure 4C**). Effector probe signal was sufficiently specific that we were able to assign dCas9 effectors at single-cell resolution by associating each individual probe to its associated effector and mapped to expected library-encoded effector combinations and set a threshold of 80% of total effector UMIs in a cell to an expected combination. To assign sgRNAs we applied the same mixture model as in benchmarking experiments above^18^.

To confirm that our experiment and perturbation assignments were successful, we first examined gene expression in a subset of cells that were assigned a strong, repressive control-dCas9-KRAB effector (“CRISPRi”) or a strong, activating VP160-dCas9-MSX2 effector (“CRISPRa”). Specifically, we looked at the impact of these effectors on the expression of CD55, which is expressed in wild-type K562 cells, and on the expression of CXCR4, which is not expressed. As expected, control-dCas9-KRAB reduced the expression of CD55, but not CXCR4, in cells containing sgRNAs targeting the promoters of CD55 and CXCR4 respectively (**Figure 4D**). In contrast, VP160-dCas9-MSX2 increased the expression of each of these genes in cells with cognate sgRNAs (**Figure 4D**).

We then expanded our analysis of cells expressing sgRNAs targeting CD55 and CXCR4 to all 19 dCas9 effectors (**Figure 4E**). For CD55, we observed robust knockdown with control-dCas9-KRAB (96.6% repression) and KRAB-dCas9-DNMT3A-3L (97.3% repression), but also substantial transcriptional repression with less characterized repressive fusion proteins such as MBD2-dCas9-PRDM2 and SaII-dCas9-CobB. Similarly, for CD55 and CXCR4, we measured robust induction of transcription, in particular with HSF1-, VP64- and VP160-containing effectors (**Figure 4E-F**). The impact of various dCas9 effectors on CD55 RNA expression measured by ProPer-seq correlated strongly with previously reported effect on CD55 protein expression measured with the COMET platform by a FACS effector screen (r = 0.9, p-value < 0.01, **Figure 4G**)^21^.

We next wanted to measure whether ProPer-seq library-on-library experiments would provide sufficient sensitivity to detect dCas9 effector regulation of target genes at the level of both specificity and secondary effects induced by gene activation. As expected, we found no impact on gene expression of a sgRNA targeting CD55 in pseudobulked cells expressing control dCas9 effector (**Figure 4H**). By contrast, we were able to detect activation of CD55 by HSF1-dCas9-FOXA1 effector, repression of CD55 by control-dCas9-KRAB effector, and activation of CXCR4 by VP160-dCas9-MSX2 effector without significant secondary transcriptional changes, which is the expected result for these genes (**Figure 4I-K**). Finally, we detected both target gene and biologically expected secondary transcriptional changes upon activation of MYOD1, a transcription factor with major roles in muscle differentiation, by VP160-dCas9-MSX2 effector (**Figure 4L-M**).

### CoMPoSE based highly multiplexed perturbation profiling

Modularity and redundancy of biological systems yield themselves to highly multiplexed perturbation workflows. For example, induction of pluripotency in somatic cells requires overexpression of at least three transcription factors, implying that unbiased discovery of novel differentiation regulatory networks would benefit from the ability to activate multiple genes at once in a parallel fashion. Conversely, gene cis-regulatory elements (CREs) form “modules” that typically only act on genes within ∼100kb distance. This makes it possible to perturb individual CREs at multiple separate loci in a single cell without inducing epistatic effects, reducing the cost of such experiments. We investigated whether our ProPer-seq workflow could be applied to the latter highly multiplexed design by perturbing multiple regulatory elements per cell in a pooled fashion in conjunction with an effector library.

Specifically, we cloned a single-sgRNA lentiviral library with 142 sgRNAs, including 12 negative control, 54 promoter-targeting ^22^ and 76 distal CRE-targeting sgRNAs^9,23–25^, co-expressing BFP. We also cloned a lentiviral library with 25 dCas9 effector constructs co-expressing mNeonGreen. We infected K562 cells first with the sgRNA library at a high MOI and then with the dCas9 effector library, resulting in 3,550 unique combinations. Nine days after the initial infection we flow sorted cells expressing both the effector and gRNAs (BFP+/mNeonGreen+) and performed ProPer-seq using custom designed effector and gRNA probes (**Figure 4A-B**).

After quality filtering, we retained a total of 303,612 single cells, 69% of which were assigned an effector and at least one sgRNA (**Figure 5A**). Supporting this assignment, cells without confident effector assignment had lower expression of mNeonGreen and dCas9 while expressing similar BFP and total RNA counts (**Extended Data Fig. 5A**). We found that 88% of our probe UMIs were contained in cells with the corresponding effector assignment, indicating high specificity of this approach. (**Extended Data Fig. 5B**). Cells had 3.5 sgRNA assignments on average, as expected from the high MOI infection, with high sgRNA UMI coverage per cell (**Extended Data Fig. 5C**, mean sgRNA UMIs per cell: 254, mean UMIs per sgRNA per cell: 72). This compression of the library-on-library experiment enabled us to robustly sample thousands of dCas9 effector-sgRNA combinations at a mean representation of 180 cells per combination (**Extended Data Fig. 5D**).

**Figure 5:**
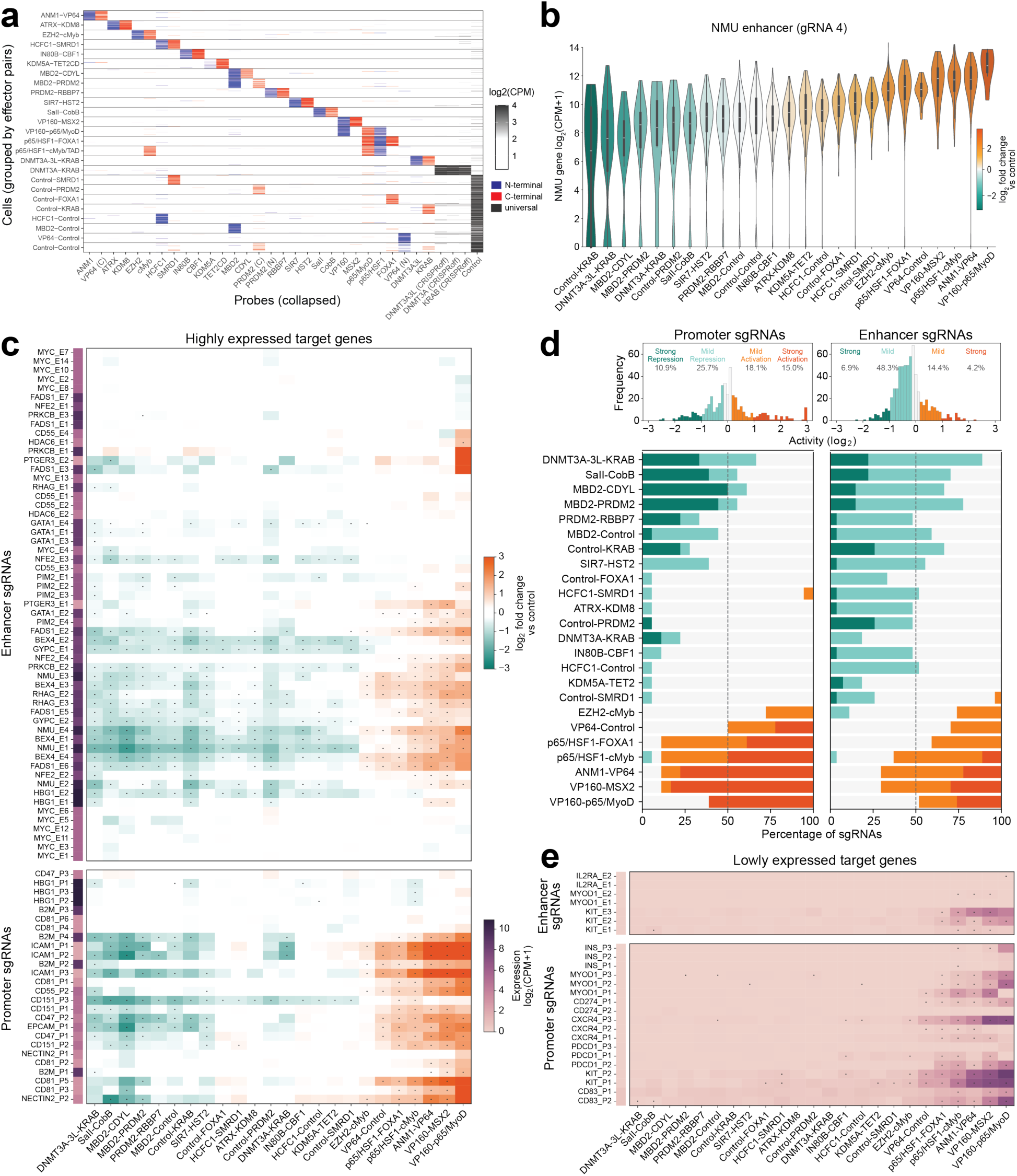
CoMPoSE based highly multiplexed perturbation profiling. **a.** Normalized count of effector probe UMIs per cell. Probes targeting the same effector are col-lapsed to the maximum of normalized probe counts. Each effector pair is represented by 100 randomly selected cells. **b.** Impact of 25 dCas9 effectors on NMU expression in individual cells expressing an NMU en-hancer targeting sgRNA compared to the average of control-effector cells containing the same NMU enhancer targeting sgRNA. Effectors are ordered by activity and plotted with shade pro-portionate to average log_2_ fold-change between that effector and control. **c.** Impact of 83 sgRNA x 24 dCas9 effectors on target gene activity of expressed genes (CPM>1 in cells with control effector) compared to control effector. Effectors are ordered by av-erage activity and sgRNAs are hierarchically clustered using correlation distance. Side annota-tion indicates basal expression level of target genes in control effector cells. Significant effector phenotypes are denoted by a point (FDR-adjusted p < 0.05). **d.** Histogram showing the distribution of target gene activity measurements for 24 effector x 45 sgRNA combinations (N = 1,080). Only sgRNAs that had a significant impact with at least 3/24 effectors are included. Activity is stratified into five functional categories: neutral (−0.2 < log2FC < 0.2), mild activation (0.2 < log2FC < 1), strong activation (log2FC > 1), mild repression (−1 < log2FC < −0.2), and strong repression (log2FC < −1). Stacked bar chart displays the percentage of sgRNAs (out of 45) exhibiting each regulatory class for individual dCas9 effectors, ordered by mean effector activity. **e.** Effector by sgRNA heatmap as in (**c**) shown for lowly or unexpressed target genes (CPM<1, N = 25 sgRNAs, 24 effectors).

To demonstrate that the presence of multiple sgRNAs per cell need not substantially affect the outcomes, we compared the impact of targeting different dCas9 effectors to the CD55 promoter in high MOI context with results obtained previously at low sgRNA MOI. We found that these results were highly correlated (Pearson’s r = 0.88, p-value < 0.01, **Extended Data Fig. 5E**). We also confirmed that targeting different dCas9 effectors to enhancers had similar average impact on target gene expression as promoter targeting (**Figure 5B, Extended Data Fig. 5F-G**). Effectors typically maintained a repressive or activating function across a range of loci (**Figure 5C-D; Extended Data Fig. 5H**). While most sgRNAs directed similarly strong impact on gene expression whether complexed with repressive or activating effectors, some sgRNAs consistently failed to affect their target genes, highlighting the need for sgRNA selection in epigenomic editing context (**Figure 5C**). Finally, we demonstrated that some of the strongest activating effectors, containing VP160, VP64 and HSF1 domains, were able to induce expression of a range of target genes with very low expression in wild-type K562 cells, including c-Kit, CD274/PD-L1, CD83 and CXCR4 (**Figure 5E**).

Together, this work demonstrates that single cell multiplexed combinatorial perturbation experiments enabled by CoMPoSE can interrogate modularly engineered dCas9 effectors across genomic contexts. More broadly, CoMPoSE is a method for the evaluation of modular synthetic effectors across genes with broad application in cellular and genomic engineering.

### A CoMPoSE library-on-library approach for scalable primary T cell engineering

The ability to systematically profile combinatorial synthetic protein designs in human cells remains a bottleneck for therapeutic engineering. CAR T cells exemplify this challenge. While curative for specific blood cancers, broader application against solid tumors has been challenging, primarily due to T cell exhaustion and immunosuppressive tumor microenvironments^26^. Multiple engineering approaches are being pursued to overcome these limitations, including modification of CAR intracellular signaling domains, genome editing of T cells and overexpression of transgenes such as transcription factors^6,27–31^. Understanding the impact of different combinations of synthetic interventions on a wide variety of T cell functional states requires a highly multiplexed single cell approach.

To systematically dissect how CAR structure and transcriptional regulators combine to control T cell functional states, we designed a library-on-library CoMPoSE experiment leveraging both the custom probe capabilities and fixed cell workflows of the ProPer-seq single cell profiling approach (**Figure 6A**). We generated a lentiviral library encoding 26 anti-CD19 CARs (GFP-tagged) with diverse costimulatory and signaling domains, and another lentiviral library encoding 10 ORFs (mCherry-tagged) including 9 transcription factors and signaling proteins previously shown to modulate T cell function^27,31^ and a control. Co-infection of activated primary human T cells from two donors with both libraries generated 260 experimental CAR × ORF combinations. We co-cultured this pool of engineered CAR-T cells with CD19-expressing A375 target cancer cells in a repetitive stimulation challenge, replacing A375 cells every two days, to model CAR T cell activation and exhaustion^28^. At the third stimulation, we split CAR T cell samples into IL-2 supplemented and withdrawn conditions. We fixed cell aliquots under three conditions - at the second stimulation, the fifth stimulation with IL-2 and the fifth stimulation without IL-2. All fixed samples were sorted for presence of a CAR and an ORF (GFP+/mCherry+) by FACS. CoMPoSE’s fixed-cell workflow enabled probe hybridization and processing of all timepoints simultaneously, minimizing batch artifacts. Custom probe panels tiling CAR and ORF transgenes enabled single-cell identity assignment across the combinatorial space (**Figure 6A**).

**Figure 6:**
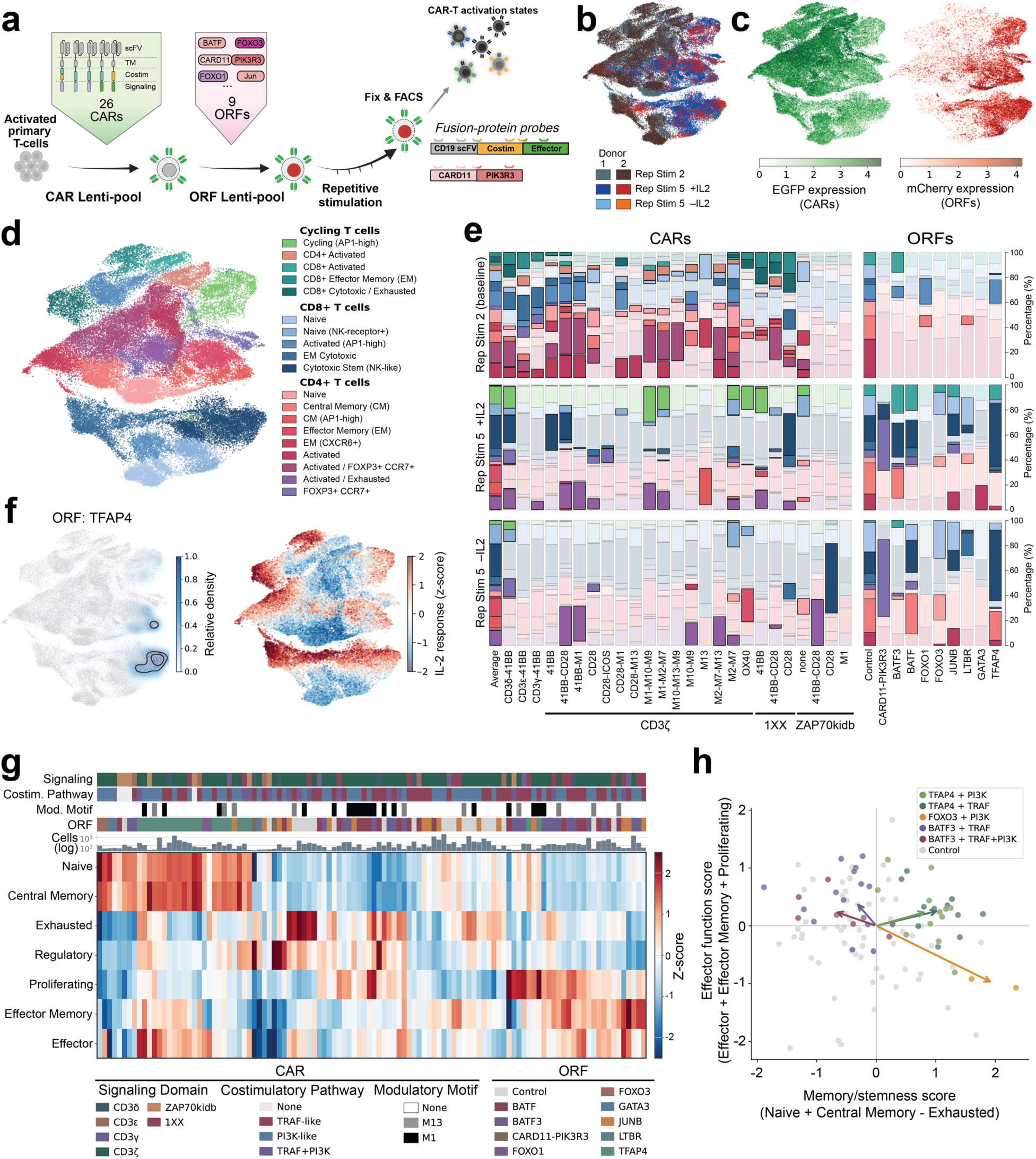
A CoMPoSE library-on-library approach for scalable primary T cell engineering. **a.** Experimental workflow for a CAR and ORF CoMPoSE library-on-library experiment in T cells. The CAR and ORF lentiviral libraries were infected serially into activated primary T-cells which were challenged with a repetitive antigen stimulation assay. Cells were sampled after 2- and 5-48-hour coculture repetitive stimulations and fixed. Fixed cells were then sorted for CAR and ORF expression based on fluorescent protein markers co-encoded on the lentiviral transgenes. These sorted cells were then processed with CoMPoSE to generate transcriptional T-cell phe-notypes and assign transgene identity. **b.** Distribution of single cells by donor (N = 27,337 from donor 1, 42,463 cells from donor 2) and treatment (N = 42,050 cells in second repetitive stimulation, 21,784 cells in fifth repetitive stimu-lation with IL-2, and 5,966 cells without) in UMAP space. **c.** CAR (EGFP probe) and ORF (mCherry probe) expression in UMAP space. **d.** Cell type annotations. See markers levels in **Extended Data** Fig. 6a. **e.** Cell type distribution for each CAR and ORF. CARs are sorted by signaling CD3ζ, CD3ε, CD3δ, CD3γ, or 1XX, ZAP70_kidb_ domains. Significantly depleted or enriched cell types compared to average distribution (CARs) or to distribution in cells containing control ORF (ORFs) are shown as a solid shade (FDR-corrected Fisher’s exact p < 0.05). **f.** Left: Location of T cells assigned TFAP4, as a representative ORF perturbation, in UMAP space. Right: Pathway activity z-score for IL-2 response (GSE39110) in UMAP space. **g.** Average impact of CAR and ORF combinations in fifth repetitive stimulation with IL-2 showing on phenotype signature z-scores (Naïve, Central Memory, Effector Memory, Effector, Ex-hausted, Proliferating, and Regulatory). CAR domains are annotated by signaling domain, cost-imulatory pathway or modulatory motifs. **h.** Scatter plot of memory/stemness pathway activity and effector function across CAR and ORF perturbations. Each CAR x ORF perturbation grouped by costimulatory domain family is plotted with the donor replicate average effector function score (effector + effector memory + prolifera-tion) and memory/stemness score (Naïve + central memory - exhausted) phenotype z-scores. Vectors are the average change in phenotype for each CAR x ORF combination.

High quality transcriptomes obtained from 69,800 T cells across two donors and three conditions showed clear separation by treatment condition (**Figure 6B**). ORF expression increased throughout the repetitive stimulation challenge (5 to 31 mean UMIs from second to fifth stimulation with IL-2), reflecting selection for ORF expressing T cells, while CAR expression declined (120 to 27 mean UMIs), consistent with activation-induced receptor downregulation (**Figure 6C**). Cells spanned diverse T cell functional states including naive, memory, effector, and exhausted populations (**Figure 6D**, **Extended Data Fig. 6A**). Further, CAR expression level correlated positively with transcriptomic markers of exhaustion, activation, and cytokine signaling and negatively with memory markers under all conditions. At second stimulation, we also noted a negative correlation between CAR expression and TCR signaling markers (**Extended Data Fig. 6B-D)**.

Between the second and fifth repetitive stimulation challenge, T cells shifted from naive to activated and effector memory states, with IL-2 withdrawal dramatically reducing cycling populations, as expected (**Extended Data Fig. 6D**). At the early, second stimulation, CAR architectural features drove substantial variation in lineage distributions (**Figure 6E**, left). Notably, CARs with alternative signaling domains — including CD3δ, CD3ε, CD3γ, and a single-ITAM CD3ζ variant (1XX) — significantly increased cycling and activated populations compared to conventional CD3ζ designs. These early CAR-driven effects suggested that signaling architecture shapes initial T cell responses to antigen encounter.

In contrast to the early effect of CARs, ORFs had a much stronger impact later, at the fifth repetitive stimulation (**Figure 6E**, right). Each ORF imposed a distinct phenotypic program consistent with its known biology (**Figure 6F, Extended Data Fig. 7A-C**)^31^. For example, TFAP4, BATF and BATF3 induced overlapping programs characterized by increased activation markers, chemokine receptors (CCR7, CCR8), and cytotoxic effector genes (GZMB, GZMK), FOXO1 and FOXO3 induced memory-associated signatures with elevated expression of stemness factors (TCF7, LEF1, IL7R), while CARD11-PIK3R3-expressing cells were enriched in a CD4+ cluster characterized by increased expression of FOXP3, CCR7, IL-2RA and checkpoint receptors (TIGIT, CTLA4, LAG3) and lower expression of memory regulators (TCF7, LEF1).

To systematically evaluate the relative contributions of CAR architecture and ORF identity across the full combinatorial space, we analyzed activation and differentiation signatures in all 260 CAR-ORF combinations at fifth stimulation, supplemented by IL-2 (**Figure 6G**). Unsupervised hierarchical clustering of these signatures revealed that ORF identity was the primary organizing principle, with distinct transcriptional programs maintained across diverse CAR architectural variants. We further visualized this hierarchical relationship by projecting all combinations into a two-dimensional space defined by memory/stemness (Naive + Central Memory - Exhausted signatures) and effector function (Effector - Effector Memory + Proliferating signatures), clearly demonstrating the dominant impact of ORFs (**Figure 6H**). In conclusion, we demonstrated that CoMPoSE enabled exploration of a highly combinatorial space (260 transgene combinations) in primary human cells under multiple conditions, which would be highly challenging with current standard perturb-seq approaches.

## DISCUSSION

Single-cell functional genomic techniques like perturb-seq are powerful approaches to scalably and richly phenotype gene function across contexts. Substantial recent attention has been driven by the application of these approaches in concert with predictive machine learning approaches to model and anticipate gene function across contexts. A requirement of such efforts is high quality perturbational data that efficiently and robustly captures large numbers of perturbation phenotypes. Here, we have demonstrated that ProPer-seq, a custom DNA probe-based single-cell transcriptomic profiling approach has key features including high scalability, fixed cell profiling, robust detection of synthetic genes, significantly reduced cost, and increased perturbational resolution in single cells. These qualities resulted in ProPer-seq being selected as the method for generating training data for the Virtual Cell Competition^3,16^. Our data also establishes that we can use ProPer-seq to validate machine learning predictions of the cellular consequences of genetic perturbation. In summary, ProPer-seq enables efforts to train and validate deep learning models that can accurately predict phenotypes arising from genetic perturbations^2,3,16,20^.

Beyond standard perturb-seq approaches, we then demonstrate that CoMPoSE library-on-library experiments using synthetically engineered proteins can be used at single cell resolution to search for optimal solutions for programming transcription or for programming cell function. We conclude that ProPer-seq is a generalizable and robust assay for rich single-cell phenotyping of complex multiplexed perturbations, including in primary T cells. Our work presents specific use cases of ProPer-seq which together serve to demonstrate that this approach is likely broadly applicable to complexly engineered synthetic proteins and combinatorial gene perturbations; and could enable multiple new types of combinatorial single cell experiments to richly characterize the perturbation effects of synthetic effectors in programming cell function.

## DATA AVAILABILITY

Raw and processed sequencing data will be made accessible on Gene Expression Omnibus upon final publication. Raw data is hosted at: SRA SUB15796302.

For reviewers, the processed data is hosted at: https://www.dropbox.com/scl/fo/fmchyizp2htuxnhmz7dwf/AJfLmlxVF5NRBwZUZoZsTy4?rlkey=c08lywegq4ent11vkmlpu2t3b&st=o6khfdma&dl=0

## CODE AVAILABILITY

CellRanger 9.0 is available from 10x Genomics https://support.10xgenomics.com/single-cell-gene-expression/software/downloads/latest

Python scripts and Jupyter notebooks for custom-probe design are available at https://github.com/jswinderman/custom_flex_probe_design

Python scripts for perturb-seq processing pipeline for Cellranger outputs is available at https://github.com/aidanwinters/pyturbseq.

## ACKNOWLEDGEMENTS

We thank all members of the Gilbert lab for helpful advice and support. The Marson laboratory has received research support from the Parker Institute for Cancer Immunotherapy, Chan Zuckerberg Institute/Biohub, the Emerson Collective, Arc Institute, Juno Therapeutics, Epinomics, Sanofi, GlaxoSmithKline, Gilead, and Anthem, as well as reagents from Genscript, Illumina, Ultima, and 10X. P.D.H. is supported by funding from the Arc Institute, Yosemite, the Biswas Foundation, the Rainwater Foundation, the Curci Foundation, the Rose Hill Innovators Program, S. Altman and V. and N. Khosla and by anonymous gifts to the Hsu laboratory. The remaining authors are or were supported by the Arc Institute.

## AUTHOR CONTRIBUTIONS

Research design, J.T.S and L.A.G. Writing and figure construction, J.T.S, M.K, and L.A.G with input from all authors. VCC experimental design, analysis, and generation J.T.S, P.Y.T, A.W, L.R.B, N.T, A.D, T.H, H.G, D.P.B, O.D.H, Y.H.R, S.K, N.L, and L.A.G. Cas9-fusion cloning, C.W. scRNA-seq J.T.S, P.Y.T. Data analysis J.T.S, A.W. CAR variant and ORF library cloning, J.T.S. Primary T cell culture, lentiviral infection, and sorting, J.T.S, L.G. Supervision, L.A.G, F.Y.F.

## METHODS

### Cell culture and lentiviral production

All cell culture was performed in a controlled humidified incubator at 37°C and 5% CO2. Cells were cryopreserved with 10% DMSO in FBS and stored in liquid nitrogen. All cell lines were tested for Mycoplasma (Lonza, cat. LT07-710) at 8 to 12-week intervals.

K562 (American Type Culture Collection (ATCC), CCL-243, female) were maintained in RPMI-1640 supplemented with 10% FBS (VWR), 1X penicillin/streptomycin with glutamine (Gibco). All K562 library lentiviral infections were performed via ‘spinfection’ at 1000 x g, 37°C, for 1-2 hours with complete RPMI supplemented with 8 µg/µL polybrene transection reagent (Millipore).

#### Lentiviral production

Unless otherwise specified lentivirus was produced in LentiX-293T, (Takara Bio, 632190) cultured in 3 mL DMEM supplemented with 10% FBS (VWR) and Pen-Strep (Gibco). Briefly, 400-600K cells were seeded onto 6 well plates the day before transfection. Per condition, 1.5 µg transfer vector, 1.35 µg of packaging vector pCMV-dR8.91, 165 ng of envelope vector pMD2.G were combined with 7.5 µL of TransIT-LT1 (Mirus Bio, MIR 2300) in 300uL Opti-MEM (Gibco) and added to the cells. Approximately 16 hours later, media was replaced with fresh media supplemented with 1X ViralBoost reagent (ALSTEM Bio). After 48 hours, viral supernatant was harvested, filtered using 0.45 µm PVDF filters and either used fresh or aliquoted and stored at − 80°C. For high MOI and T cell transductions lentivirus was produced with LV-MAX Lentiviral Production System (Thermo, cat. A35684) per manufacturer’s instructions. LV-MAX cells were subcultured in LV-MAX serum-free production medium after reaching > 95% viability. Suspension cells were transfected in 2 L flasks and 5-6 hours later, LV-MAX Transfection enhancer was added. After 48 hours, cultures were spun at 1300 x g for 15 minutes. Supernatant was filtered through 0.45 µm filters, processed using Lenti-X concentrator (Takara, 631232) and resuspended in complete RPMI and preserved at −80°C.

Functional titers of libraries were derived by transduction of K562 cells with serially diluted virus and flow cytometric assessment of effector (mNeonGreen) and sgRNA (BFP) fluorescence with an Attune NxT Flow Cytometer (Thermofisher, A24858) and CytKick Autosampler (Thermofisher, A42901). In the final experiments, infected cells were sorted to purity on a BD FACSAria Fusion (BD Biosciences, 657590).

#### Primary T cell isolation culture, and repetitive stimulation

Human T cells were isolated from peripheral blood mononuclear cell enriched Leukopaks from healthy donors (StemCell Technologies 200-0092) and CD3+ were isolated by EasySep Human T cell isolation kit (100-069). Unless otherwise specified T-cells were cultured in complete X-VIVO-15 (X-VIVO-15 (StemCell Technologies, 04-418Q) supplemented with 5% fetal calf serum (R&D systems, lot M19187), 1X Pen-Strep (Gibco), and 50 IU/mL IL-2 (R&D systems, 202-GMP-01M). T-cells were cultured at 1 x 10^6^ cells/mL in complete X-VIVO-15 and activated with anti-CD3/CD28 Dynabeads (Life Technologies, 40203D) at a density of 1 x 10^6^ cells/mL. After 48 hours of activation T cells were separated from Dynabeads with a magnetic tube column and ‘spinfected’ at 1000 x g for 1 hour at 37°C with 2.5% v/v of 100x Lenti-X concentrated CAR library lentiviral pool in fresh complete X-VIVO-15. After 16 hours the same cells were then ‘spinfected’ at 1000 x g for 1 hour at 37°C with 5% v/v of 100x Lenti-X concentrated ORF library lentiviral pool and resuspended at 1 x 10^6^ viable cells/mL.

Repetitive stimulations were conducted with CD19^+^ nuclear RFP^+^ A375 melanoma cells which were cultured and seeded in complete RPMI. Complete RPMI was removed, and T cells were plated on A375 target cells at 1:1 ratio in complete X-VIVO-15 supplemented with fresh IL-2 and cultured for 48 hours. After 48 hours CAR ^+^ and ORF^+^ were tracked by Attune NXT Cytometer (Invitrogen) and cell viability by Countess. T cells were resuspended in freshly prepared complete X-VIVO-15 with IL-2 and plated on fresh A375 target cells. This was repeated for 5 repetitive stimulations. At repetitive stimulation three each donor was resuspended in divided evenly and resuspended in either complete X-VIVO-15 with or without IL-2 and plated on fresh A375 target cells. At two repetitive stimulations half of the T cells were fixed and enriched by FACSAria Fusion (BD Biosciences, 657590). After repetitive stimulation five cells cultured in IL-2 complete and withdrawn media were similarly fixed overnight and sorted by FACSAria Fusion (BD Biosciences, 657590) prior to quenching and storage at −80°C (see *Sample fixation* below).

### sgRNA and effector library design and cloning

#### Single-target and GEARS nominated genetic interaction library

To construct the genetic-interaction library, GEARS was trained on K562 CRISPRa single-and dual- gene perturbations. All predicted gene-gene combinations were scored for model fit, magnitude, and similarity of single to dual (see below) fit metrics. GIs with varying fit metrics across these predicted parameters were selected while minimizing number of individual single perturbations. A programmed dual-sgRNA library was designed and ordered as oligo-pools from Twist Bioscience. sgRNA-pairs were represented scaled to previously measured growth fitness effects^19^. sgRNA scaling values are provided in **Extended Data Table 1** and shown in **Extended Data Fig. 2C** (scaling factor estimated as 2^-γ*4^, assuming essential perturbations begin depleting ∼48 hours after sgRNA lentiviral transduction: γ is a previously defined doubling normalized fitness term^18^. Pooled sgRNA libraries were quality controlled by Whole Plasmid Sequencing (Plasmidsaurus), and NGS of the plasmid library with a Miseq V3 (Illumina).

#### Low-, high- MOI library and dCas9-effector plasmid cloning

For high-MOI libraries protospacers were curated from the literature as detailed in **Extended Data Table 1**. When necessary, protospacer sequences were trimmed to the 19- PAM proximal bases and the 20th most PAM-distal nucleotide was mutated to a ‘G’. Single-guide RNAs were ordered as DNA oligo-pools from Twist bioscience equimolarly. They were digested and ligated into a lentiviral sgRNA expression vector pJR103 via BstXI and BlpI. Pooled sgRNA libraries were quality controlled by Whole Plasmid Sequencing (Plasmidsaurus), and NGS of the plasmid library with a Miseq V3 (Illumina).

Arrayed COMET effectors were cloned into a lentiviral vector containing SFFV-XTEN16-2XNLS -dCas9-XTEN80 (pCW4)^21^ by PCR-amplifying synthesized effector gene fragments (Twist Bioscience) and assembled by Gibson-cloning (2X HiFi Assembly Master Mix, NEB).

#### CAR variant and ORF plasmid cloning

To create a lentiviral backbone adaptable to scaled golden-gate assembly insertion of variable signaling and costimulatory domains pZR135^32^, an anti-CD19 scFV-CD8a hinge-CD28-CD3ζ-T2A-EGFP backbone was used as template to introduce tandem PaqCI sites 5’ of the CD8a hinge and 3’ of the T2A site. ORF fragments were then designed to reconstruct in-frame costimulatory and signaling domains 3’ of the CD8a hinge and 5’ of the T2A-EGFP. Recoded costimulatory and signaling domains were ordered as Gene Fragments (TWIST Biosciences, ordered fragments provided in **Extended Data Table 3A**). Golden Gate Assembly reactions with PaqCI and T4 ligase (New England Biolabs, R0745 and M0202) were set up for each CAR variant; each insert was mixed at a 2:1 molar ratio with 50 ng destination plasmid, 1 µL 10X T4 DNA Ligase Buffer, 0.5 µL 10 u/µL PaqCI, 0.25 µL 20 µM PaqCI Activator, and 400 u/µL 0.5 µL T4 DNA Ligase to a final reaction volume of 10 µL with nuclease-free water (Ambion). Reactions were incubated at 37°C for 60 minutes, and 0.5 µL transformed into 10 µL of Stellar *E. Coli* chemically competent cells (Takara). Colonies were screened for correct insertion size by Colony PCR and transformants with correct insert size were then selected for final verification by Whole Plasmid Sequencing (Plasmidsaurus).

An E1fα promoter was cloned 5’ of a recoded ORFs followed by P2A and mCherry by PaqCI Golden Gate Assembly as above. The E1fα promoter, ORF fragments and P2A-mCherry were ordered as Gene Fragments (TWIST Biosciences, ordered fragments provided in **Extended Data Table 3B**). Golden Gate Assembly reactions with PaqCI and T4 ligase (New England Biolabs, R0745 and M0202) were set up for each ORF variant; 1-3 ORF inserts with E1fα and mCherry were mixed at 2:1 molar ratio with 75 ng destination plasmid, 1.5 µL 10X T4 DNA Ligase Buffer, 0.75 µL 10 u/µL PaqCI, 0.375 µL 20 µM PaqCI Activator, and 400 u/µL 0.75 µL T4 DNA Ligase to a final reaction volume of 15 µL with nuclease-free water (Ambion). Reactions were incubated at 37°C for 60 minutes, and 0.5 µL transformed into 10 µL of Stellar *E. Coli* chemically competent cells (Takara). Colonies were screened for correct insertion size by Colony PCR and transformants with correct insert size were then selected for final verification by Whole Plasmid Sequencing (Plasmidsaurus).

### Single cell RNA-seq library preparation

#### Sample fixation and hybridization

Cell suspensions were fixed with Chromium Next GEM Single-Cell Fixed RNA Sample preparation kit (10x Genomics, 1000414) according to the Fixation of Cells and Nuclei Demonstrated Protocol # CG000478 (Rev D). Briefly, aliquots of 2*10^6^ cells were fixed in Fixation Buffer for 20 hours at 4°C, quenched and supplemented with Enhancer to 10% and 50% w/v Glycerol (Fisher Scientific, 329016) to 10% and stored at −80°C until post-fixation processing.

For the CAR x ORF experiments T cells that were harvested at RS2 or RS5 were each fixed in Fixation Solution overnight at 4°C. Fixed cells were quenched in Quench Buffer immediately before sorting, washed and resuspended in sorting buffer (PBS supplemented with 1% nuclease-free BSA and 0.2 U/µL Protector RNase inhibitor (Sigma)). Cells were sorted into sorting buffer, and resuspended in Quench Buffer supplemented with 10% Enhancer and 10% Glycerol (Fisher Scientific) prior to long-term storage at −80°C.

Cryopreserved fixed samples were thawed and ∼500,000 cells were hybridized using Chromium Fixed RNA Kit, Human Transcriptome, 4rxn x 4BC (PN-1000475, 10x Genomics) or Chromium Fixed RNA Kit, Human Transcriptome, 16rxn x 16BC (PN-1000475, 10x Genomics), LHS and RHS custom probes were ordered as oligo pools (IDT) and added to hybridization reactions according to the WTA barcode to a final concentration of 5 nM each following 10x Genomics Demonstrated Protocol #CG000621 (Rev D). For sgRNA probes the RHS sgRNA probe was included to a final concentration of 80 nM and the LHS protospacer probe was added to a final concentration of 20 nM. For all ProPer-seq experiments other than those in **Figure 1A** modified hybridization amplification protocol was applied: prior to a 16–24-hour 42°C incubation samples were ramped from 65°C at 1°C /5 minutes. Hybridized samples were pooled equally and washed following the Pooled Wash Workflow #CG000527. Two lanes of GEMs were generated on the Chromium X (10x Genomics) targeting 60,000 cells recovered.

#### Transcriptome, sgRNA, and effector library construction

For hybridized sgRNA probe amplification pre-amplification was performed with Feature Pre-Amp Primers and amp mix using Fixed RNA Feature Barcode Kit 16 rxns (10x Genomics, PN-1000419). For experiments with Read 2 Nextera effector probe pools a Nextera Read 2 complementary primer (5’-GTGACTGGAGTTCAGACGTGTG-3’) was added to the pre-amplification mixture to a final concentration 1 µM. Pre-amplification libraries were prepared according to manufacturer specification. 20 µL of this product was used as input to construct Read 2 Nextera and/or Read 2 TruSeq pre-indexing intermediate libraries. Forward and Reverse primers for Read 1 TruSeq and Read 2 Nextera or TruSeq were added at 1 µM in1X amp mix to 20 µL of pre-amplification product to construct the sgRNA preamplification product (TruSeq R2) or 20 µL of pre-amplification for effector pre-amplification (Nextera R2). Reactions were cycled according to pre-amplification cycling conditions with two additional cycles in a ProFlex PCR System (ThermoFisher Scientific, cat. A30754). These libraries were size-selected by SPRI-selection (1X; Beckman Coulter B23318) and then 20 µL was used as input for indexing with Dual Index Kit NT Set A 96 rxns (10x Genomics, PN-1000250) or Dual Index Kit TT Set A 96 rxns (10x Genomics, PN-1000215). Indexed libraries were quantified by Qubit dsDNA Quantitation, High-Sensitivity kit (Thermo Fisher Q32851) for yield and High-Sensitivity D1000 TapeStation (Agilent 5067-5585) to ensure expected library size and absence of contaminating products (representative traces shown in **Extended Data Fig. 4E-G**). Pooled libraries were paired-end sequenced: Read 1: 28, I1: 10, I2: 10, Read 2: 90 (Illumina Nova-Seq X).

## 5’ and 3’ direct-capture perturb-seq

5’ direct-capture perturb-seq libraries were prepared with freshly harvested transduced and sorted CRISPRa K562 according to manufacturer’s recommendation (10X Genomics, User Guide CG000735 (Rev A). We sought to load 26,000 cells each across 6 separate GEM-X lanes. A workflow modification was made to amplify the sgRNA in position A. A separate Feature PCR Mix was performed with 1 µM each of CR3_SI_F “5’-GATCTACACTCTTTCCCTACACGACGC-3’” and CR3_SI_R “5’-GTGACTGGAGTTCAGACGTGTGCTCTTCCGATCTCAAGTTGTAAACGGACTAGCCTTATTT CAAC-3’”. These were then separately indexed with unique Dual Index Plate TT Set A (10x Genomics PN-3000431) indices and pooled equally with the standard direct-capture sgRNA product for NGS sequencing.

3’ direct-capture perturb-seq libraries were prepared as above with K562 transduced with sgRNA libraries cloned into pJR85 lentiviral backbone and pJR89 U6-CS1 insert. Cells were processed in parallel to cells transduced with sgRNA libraries with unmodified constant region vectors and processed according to manufacturer’s recommendation (10X Genomics, User Guide CG000418 (Rev D)).

### Analysis

#### scRNA-seq analysis

BCL Convert Software (Illumina) was used to generate demultiplexed FASTQ sequencing libraries which were aligned with Cellranger multi (v.9.0.0, 10x Genomics) using a human probe-set reference (v1.0.1-GRCh38-2020-A) with CRISPR Guide Capture and corresponding sgRNA feature reference. For effector library experiments the WTA probe-set was modified to include custom probe sequence references (dCas9 effector sequences provided in **Extended Data Table 2C** and CAR x ORF sequencs in **Extended Data Table 3C**). Cell barcodes were filtered to remove cells with fewer than 1,095 UMIs or 665 genes detected, mitochondrial content greater than 15%. Only cells with two confident sgRNA assignments corresponding to expected library pairs were included.

#### Assay benchmarking

In 5’ direct-capture, 3’ direct-capture, ProPer-seq and Virtual Cell Challenge (https://www.dropbox.com/scl/fo/fmchyizp2htuxnhmz7dwf/AJfLmlxVF5NRBwZUZoZsTy4?rlkey=c08lywegq4ent11vkmlpu2t3b&st=o6khfdma&dl=0) or (https://virtualcellchallenge.org/, adata_training.h5ad) data were subsampled to 2,000 cells with sgNeg assignment and down-sampled to equal reads. and UMIs and gene per cell as well as percent of counts in top 200 genes were calculated with scanpy calculate_qc_metrics^33^. For all perturb-seq experiments on-target activity was determined as the log_2_ of target gene expression in a targeting sgRNA assigned cell over sgNeg in CPM. For a scaled ProPer-seq experiment with the Virtual Cell Challenge we recovered 844,736 cells with median 540 sgRNA UMI/cell, 93% of which had >80% on-target knockdown.

#### Genetic-interaction and perturbation analysis

Count matrices were filtered to genes with a mean UMI >0.25 and normalized to the median total counts. Gene expression was then z-score normalized to sgNeg cells, and this z-normalized matrix was pseudobulked by perturbation assignment. All pairwise correlations of these pseudobulked perturbations were then calculated with scikit-learn cosine_similarity().

We applied a robust regression linear model^19^ to decompose genetic interactions on z-score normalized perturbation pseudobulked groups to *∂ab=* c_1_ *∂a+* c_2_ *∂b+ ε* where *∂ab*, *∂a*, and *∂b* are the observed normalized expression difference of an GI pair and single compared to sgNeg pairs, c_1_ and c_2_ are fit scalars of the single perturbation phenotypes and ε is a fit error term summarizing the nonlinear interaction between ∂a and ∂b. The primary model terms described are: model fit *dcor(c_1_a+ c_2_b,ab)*, magnitude = *√( c_1_^2^+ c_2_^2^)*, and similarity of single- to dual-perturbations *dcor([a,b],ab)*. For analyses including GEARS performance evaluation the thresholds were: neomorphic as model fit < 0.88, synergistic as magnitude > 1.15, suppressive as magnitude < 1, and redundant as similarity of single to dual > 0.85.

Energy distance^10^ for all perturbations was calculated using etest against sgNeg pairs with 1000 permutations. DESeq2 was used for differential expression testing using pyDESeq2^34^ on count matrices pseudobulked by perturbation assignment compared to sgNeg.

#### Custom probe design

Custom probes to detect dCas9 effector fusions we implemented a workflow adapting 10x Genomic Technical Note CG000621 (Rev D). N- and C- terminal effectors sequences (available in **Extended Data Table 2**) were converted to FASTA from which candidate probe pairs were nominated by binning the reverse complement into 50bp fragments, splitting them into the first 25 bp (LHS probe) and last 25 bp (RHS probe) and filtering any with: GC content less than 44% or greater than 72%, homopolymer length greater than 4 bases, Shannon entropy greater than 0.8, and scored 2-5 base repeat sequences. The LHS and RHS probe for each pair was scored for off-target binding with blastn against hs.refseq.rna, and all off-target binding sites greater than 20 bases were recorded. Off-target binding length, and the sequence metrics were aggregated into a probe-pair priority score, the total priority score of all non-overlapping sets of three or fewer probe-pair sets were scored while penalizing sets less than three, the lowest scoring set was selected for each effector sequence. Probes were constructed as follows: RHS probe: “/5Phos/” + rhs_sequence + “ACGCGGTTAGCACGTANN” + barcode_sequences[i] + “CGGTCCTAGCAA” LHS probe: lhs_r2_adaptor + lhs_sequence where the lhs_r2_adaptor was “CGGAGATGTGTATAAGAGACAG” for Read 2 Nextera libraries (**Extended Data Fig. 4F**), and “CCTTGGCACCCGAGAATTCCA” for Small RNA Read 2 libraries.

For sgRNA probes design (**Extended Data Fig. 4E**), the constant sequence and probe barcode were repositioned to the LHS probe which hybridizes to the sgRNA constant region to reduce oligo pool synthesis.

Position A sgRNA RHS probe: “/5Phos/” + “CTGAACNNNNNNNNNNNNNNNNNNNG” + “CGGTCCTAGCAA”

Position B sgRNA RHS probe: “/5Phos/” + “AAACNNNNNNNNNNNNNNNNNNNG” + “CGGTCCTAGCAA”

Position A Constant Region LHS probe: “CAGACGTGTGCTCTTCCGATCT” + “ACGCGGTTAGCACGTANN” + barcode_sequences[i] + “CTTGCTATGCACTCTTGTGCTTAGCT”

Position A Constant Region LHS probe: “CAGACGTGTGCTCTTCCGATCT” + “ACGCGGTTAGCACGTANN” + barcode_sequences[i] + “GCTATGCTGTTTCCAGCTTAGCTCTT”

#### CoMPoSE dCas9 effector x CRE analysis

To assign effector terminal pairs to cells UMIs from effector probe sets were first depth normalized to UMIs (counts) per 10,000 (CP10k). For each CBC, all probe UMIs for terminal effectors were grouped by terminus and effector. Then N- and C- terminal probe UMIs were assigned as a pair if more than 80% of the total UMIs corresponded to a pair within the dCas9-effector library. An exception was made when more than 80% of UMIs were assigned to RC433 without another effector having more than 20% effector assignment. Differential expression testing was done as described above. Cells were pseudobulked according to their sgRNA and effector assignment as well as probe barcode to construct replicates. DESeq2 analysis was performed on these pseudobulks with probe barcode groups treated as replicates. Significantly upregulated genes (padj < 0.05 and log2fc > 0.5) from differential expression analysis were used as input for geneset enrichment analysis (GSEA) with GSEApy referencing pathways from KEGG_21_human GO_biological_process_2021. Minimum gene set size considered was 15, maximum 500, and 1000 permutations were performed.

For high MOI analysis, an activity matrix was constructed by calculating the on-target activity grouped by assigned effector as above. On-target activity was determined as the log_2_ fold-change in normalized counts for all cells for a given dCas9-effector with an sgRNA assignment over cells assigned that same sgRNA with Control-dCas9-Control effector assignment.

#### CoMPoSE CAR x ORF analysis

Integrated single-cell objects were processed like above. Cells were quality controlled by first identifying outliers (>5 median absolute deviations) based on mitochondrial gene percentage and percentage of counts in the top 100 genes. Counts were then normalized and log1p transformed. For UMAP visualization, dimensionality reduction was performed using principal component analysis on the top 2,000 highly variable genes. Cells were clustered using the Leiden algorithm (resolution = 0.8) on a k-nearest neighbors’ graph (k = 15, Euclidean distance). Clusters were annotated manually by marker genes identified by Wilcoxon rank-sum test with Benjamini-Hochberg correction. Manual assignments were validated by CellTypist (Immune_All_Low), which demonstrated high concordance for several identified subsets. We removed cells contained in three identified clusters associated with low-quality (272 cells) and epithelial/myeloid contaminants (194 cells).

Cell type composition was quantified as the proportion of each T cell subset within CAR variants and treatment conditions. Statistical significance was determined by Fisher’s exact test for each CAR vs all other CARs within a condition for enrichment (odds ratio > 1) or depletion (odds ratio < 1). P-values FDR-corrected by were Benjamini-Hochberg. The same procedure was conducted for each ORF compared to mCherry control.

IL-2 response z-scores were calculated using gene set enrichment scoring with Scanpy sc.tl.score_genes() with the ImmuneSigDB GSE39110 Untreated vs IL2 TREATED CD8 TCELL Day6 POST IMMUNIZATION UP pathway, MSigDB C6 v 2023.1).

Differential expression testing for ORFs was performed using Wilcoxon rank-sum test with Benjamini-Hochberg multiple hypothesis test correction (threshold adjusted p-value 0. and abs(log_2_ fold-change) > 1.

---

**Extended Data Fig 1.**
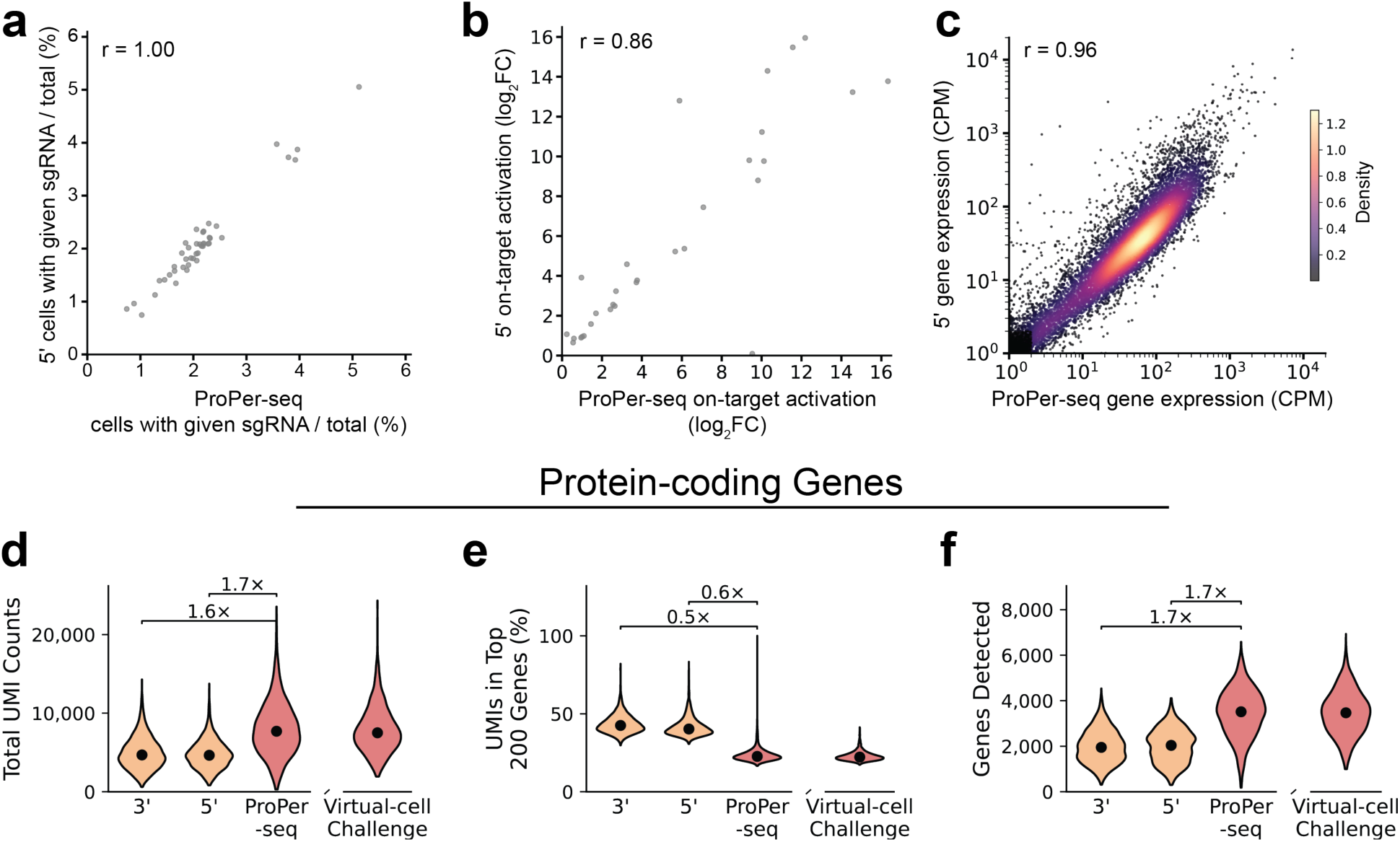
Efficient transcriptional measurement with ProPer-seq with strong perturbation phenotypes. **a.** Correlation of sgRNA representation between 5’ perturb-seq and ProPer-seq shown as fraction of cells assigned a given sgRNA out of all cells (Pearson’s r = 1.00, p<0.01, N = 42 pseudo-bulk perturbation groups). **b.** Correlation of on-target activation between the assigned guide for a single- and dual- perturbations over sgNeg against targeted genes showing ProPer-seq and 5’ perturb-seq (Pearson’s r = 0.86, p<0.01, N = 42 pseudobulk perturbation groups). **c.** Comparison of expressed gene detection level between ProPer-seq and 5’ perturb-seq, shown as density a scatter plot of log-normalized counts (UMIs)-per-million (Pearson’s r = 0.95, p-value <0.01). Genes detected below 1 CPM excluded from density. **d-f.** Single-cell violin plots of ProPer-seq and 5’ transcriptome detection showing per-cell UMIs (**d**), percent of UMIs in the top 200 most expressed genes (**e**), and total genes detected (**f**) for protein coding genes.

**Extended Data Fig 2.**
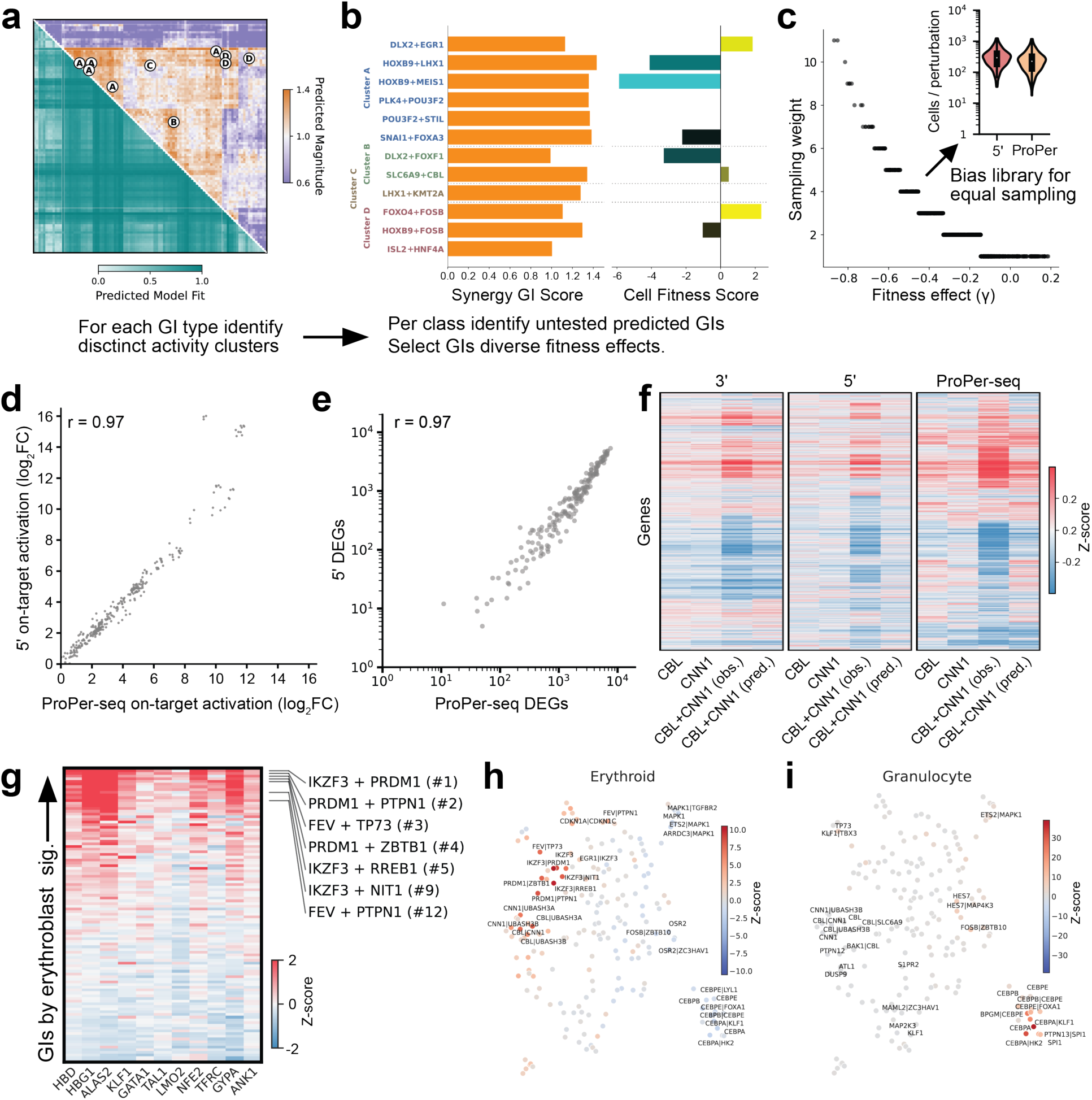
Characterization of a predicted GI phenotypes with ProPer-seq. **a.** Matrix of all 10,302 GEARS predicted GIs shown with predicted magnitude and model fit scores^19,20^. Example nominated synergistic genes are labeled. **b.** Representative GIs from various interaction class were selected with varied measured direct cell fitness effects. Example predicted synergistic GIs are shown with predicted synergy interaction scores and measured cell fitness scores^19^. **c.** To achieve balanced sampling library representation was biased based on measured fitness effect. Oligo synthesis was biased by 2^-4ˠ^ anticipating approximately four cell doublings after infection. Sampling of unique sgRNAs assigned to each cell in ProPer-seq and 5’ perturb-seq. Mean cell coverage was 259.1 ± 188.8 (ProPer-seq) and 323.5 ± 208.3 (5’ perturb-seq) single or dual sgRNA called cells. **d.** Correlation of on-target activation between the assigned guide for each single- and dual- perturbations over dual-sgNeg against targeted genes showing ProPer-seq and 5’ perturb-seq (Pearson’s r = 0.97, p<0.01). **e.** Correlation of the number of measured differentially expressed genes between each single-and dual- perturbation compared to sgNeg cells for ProPer-seq and 5’ perturb-seq (Pearson’s r = 0.97, p<0.01). **f.** Perturbation z-score matrices for CBL-CNN1 interaction measured by 3’ perturb-seq (left, 3’ reprocessed from^19^), 5’ perturb-seq (middle) and ProPer-seq (right). Plots show all linear model included consensus highly variable genes as perturbation z-scores. GIs are shown with the linear model predicted CBL + CNN1 predicted and observed transcriptional profile. **g.** Heatmap of perturbation expression z-scores of erythroblast markers for all measured GI pairs. Predicted erythroblast perturbations from the GEARS nomination are labeled with their position in the rank-order of total erythroblast marker expression across all GIs. **h-i.** Clustered pseudobulk UMAP colored by erythroid transcriptional signature (**h**), and granulocyte transcriptional signature (**i**). Applied previously defined granulocyte and erythroid signatures derived^19^.

**Extended Data Fig 3.**
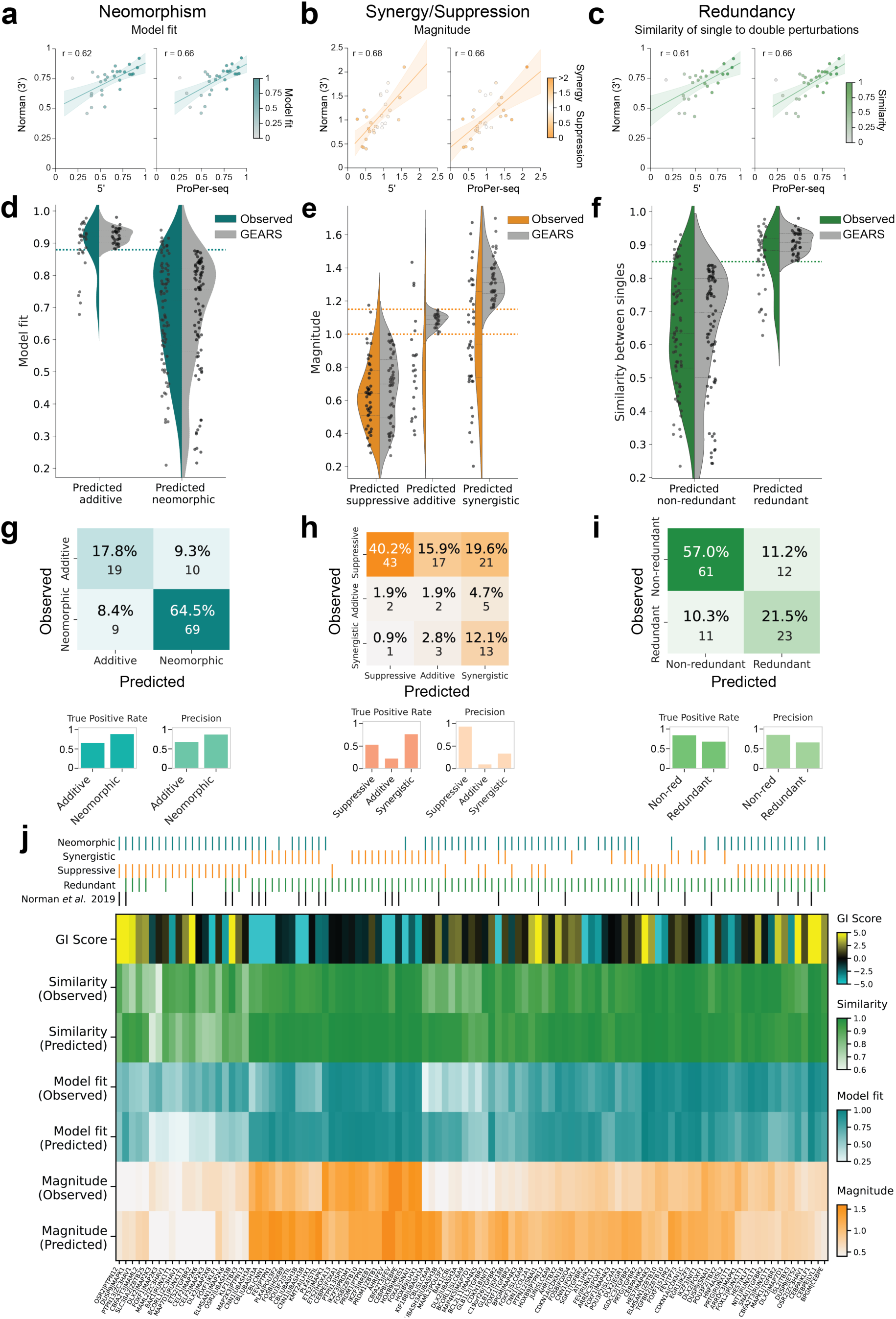
Validation of GEARS predicted genetic-interaction phenotypes. **a-c.** Correlation scatterplots between overlapping GIs from Norman *et al.*^19^ and ProPer-seq (right) and 5’ perturb-seq (left) for model fit (**a**; Pearson’s r = 0.62, and 0.66), magnitude (**b**; Pearson’s r = 0.68, and 0.66), and similarity of single to dual (**c**; Pearson’s r = 0.61, and 0.66). **d-f.** Split violin plots showing the distribution of predicted and observed terms to describe neo-morphic (**d)**, synergistic/suppressive (**e**), and redundant (**f**) genetic interactions. The threshold applied for the GEARS assignment threshold is shown. Measured distributions are shown in solid colors while the GEARS predicted distributions are shown in grey. **g-i**. Confusion matrices for the assignment precision and true-positive rate for all GEARS-predicted genetic interaction categories. **j.** Heatmap showing growth GI value measured in Norman *et al.*^19^, along with the measured and predicted similarity, model fit, and magnitude for all measured perturbations. Categorical assignments from GEARS^20^ predicted values are indicated above the heatmap along with which perturbations were replicated from the Norman *et al*. data^19^.

**Extended Data Fig 4.**
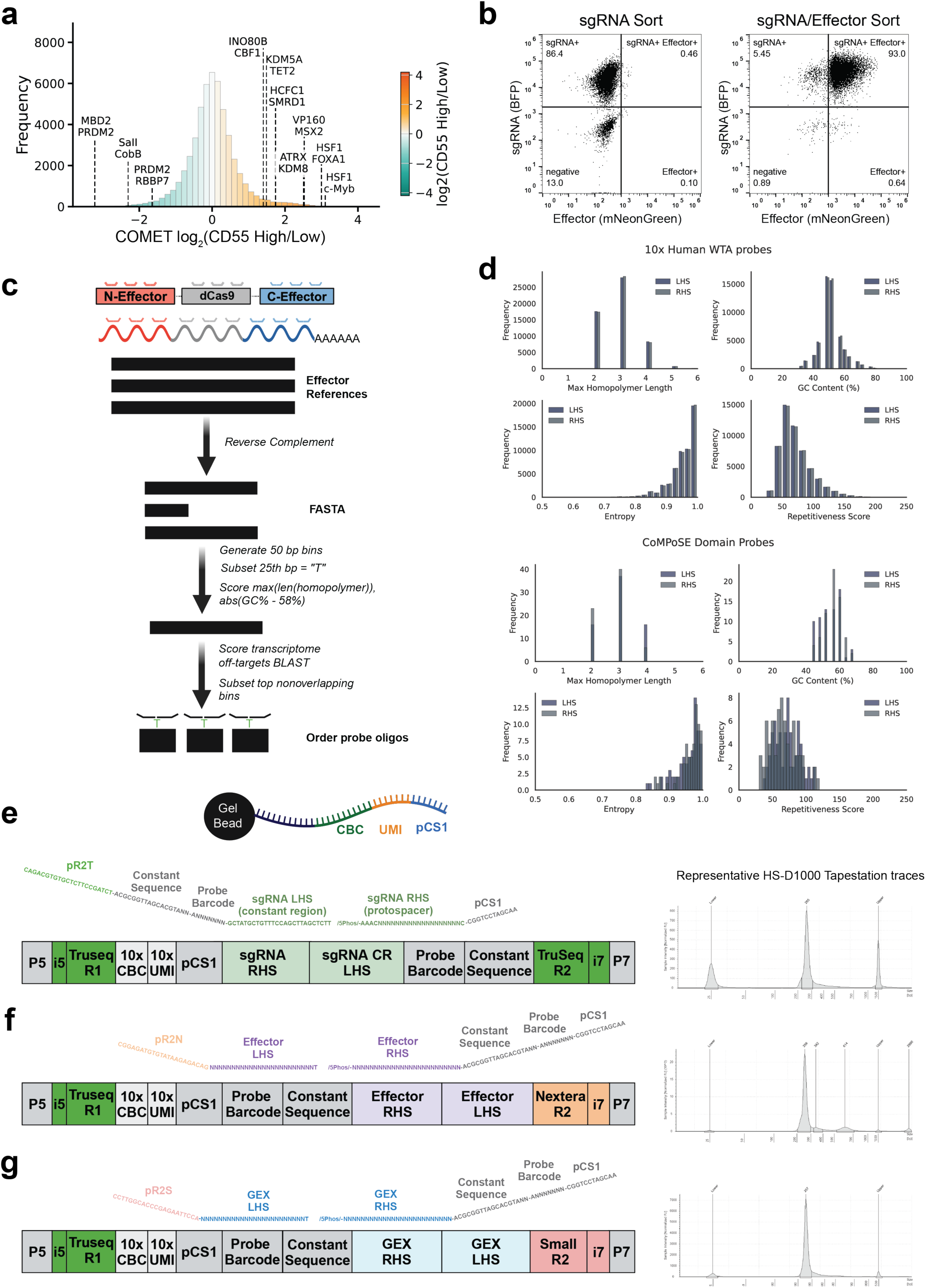
Modification of ProPer-seq for simultaneous detection of modular synthetic effectors and sgRNAs. **a.** Histogram of all observed enriched dCas9 effectors from the COMET K562 screen^21^ (https://comet-ivp.gilbertlab.arcinstitute.org/) plotted as log_2_(CD55 High/Low). Selected effectors are labeled. **b.** Cytometry for sgRNA transduced and sorted cells (left) and sgRNA and effector transduced and sorted cells (right). Plotted as log fluorescence for BFP (sgRNA marker) and mNeonGreen (effector fusion. **c.** Schematic of probe_design.py to design custom probes including binning, subsetting, annotating sequence composition, scoring off targets, and selecting optimal pair sets, **d.** Comparison of the sequence compositions; maximum homopolymer length, GC content, Shannon entropy, and repetitiveness split by LHS and RHS probe for custom probes designed for effector libraries compared to 10x produced whole-transcriptome probe-sets. **e-g.** Expected probe next-generation-sequencing library design as well as representative high-sensitivity DNA 1000 tapestation traces from NGS library construction for transcriptome (**e**), effector (**f**), and sgRNA (**g**) libraries.

**Extended Data Fig 5.**
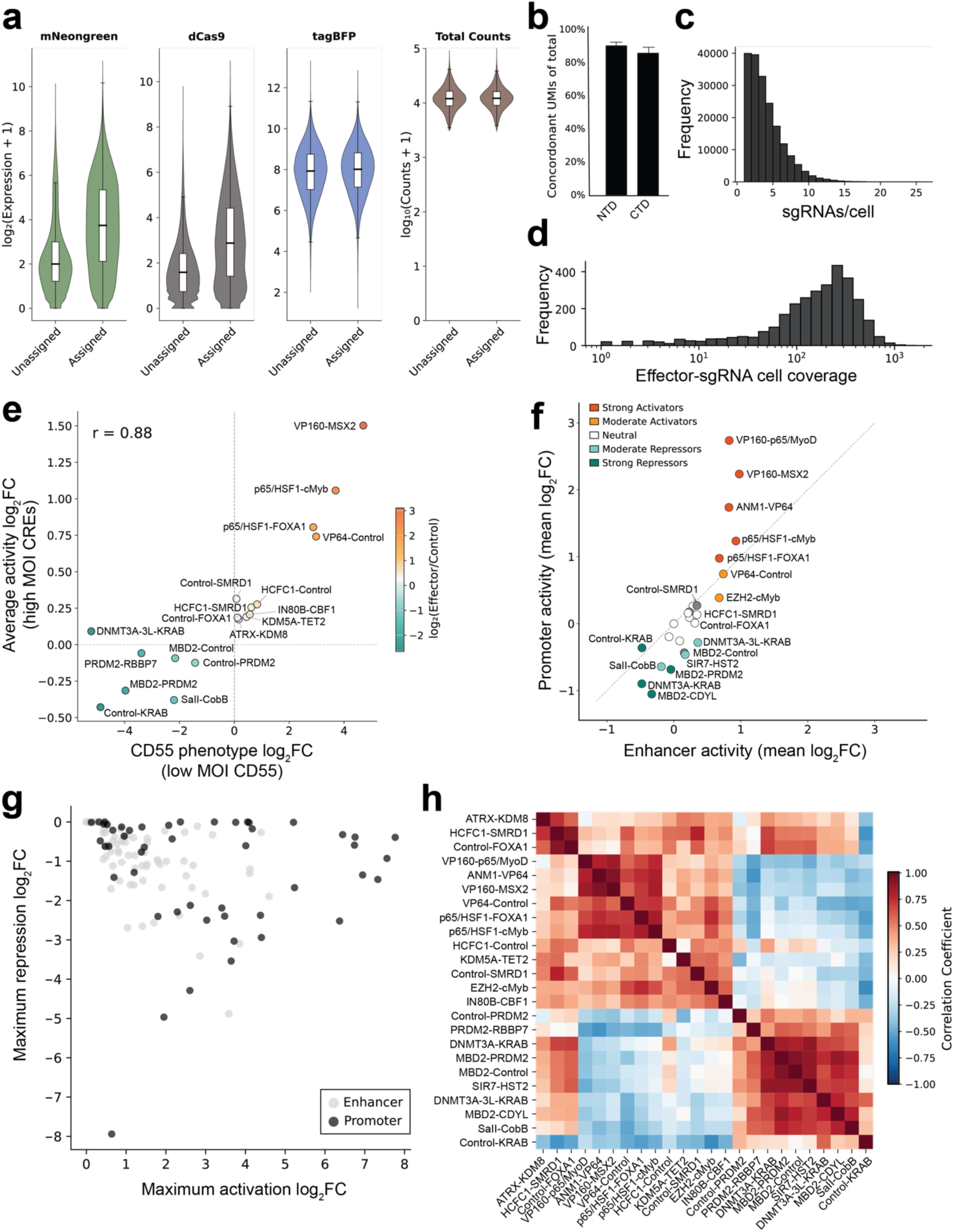
Generalized activity of dCas9 effectors across regulatory elements. **a.** Violin plot distributions of mNeonGreen, dCas9, tagBFP, and total measured reads for cells with and without confident effector assignment. **b.** Bar chart showing the sensitivity of custom N- and C- terminal domain (NTD/CTD) effector probes. The fraction of all N- and C- terminal effector probes UMIs that are consistent with the assigned effector are plotted with mean +/- SEM. **c.** Distribution of sgRNA assignments per cell (n=212,009 cells), with median of 3.5 guides per cell across the pooled library. **d.** Sampled cell distribution per sgRNA-effector combination (n=141 unique guides by 25 effectors), showing median of 3.5 guides per cell across the pooled library. **e.** Correlation scatterplot of the CD55 promoter activity measured in the CoMPoSE low-MOI experiment against the average activity across all CREs for each effector (Pearson’s r = 0.88, p-value < 0.01). **f.** Scatter plot comparing mean regulatory activity (log2 fold-change) of each effector at enhancer regions versus promoter targeting sgRNAs (Pearson’s r = 0.68, p-value < 0.01) **g.** Scatter plot of the maximally observed activation and repression strength at each CRE colored by promoter elements (black) and enhancer elements (grey). **h.** Correlation matrix of all effectors correlated on activity across all targeted CREs.

**Extended Data Fig 6.**
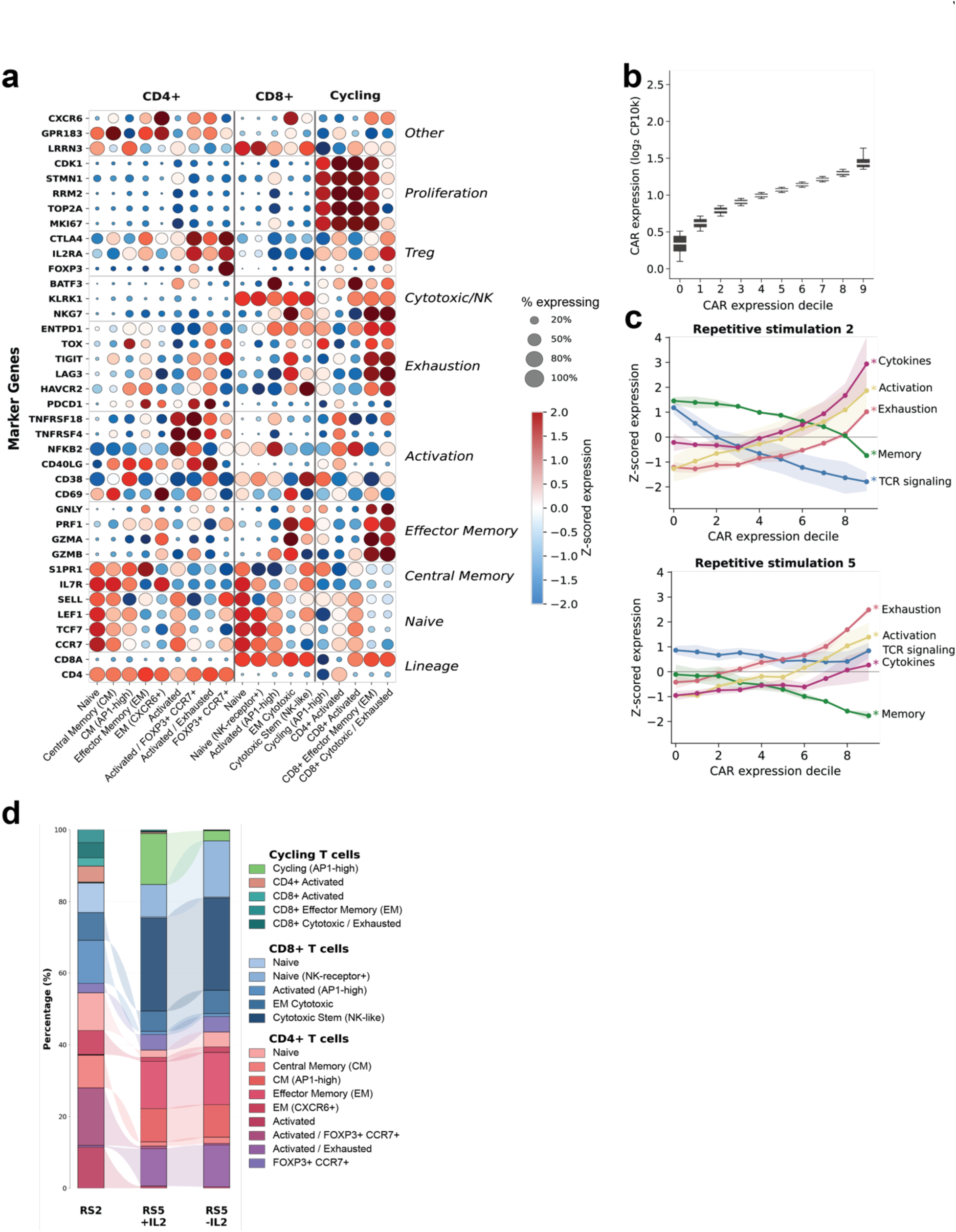
CoMPoSE measurement of T-cell activity across CAR expression and repetitive stimulation challenge. **a.** Dot plot for selected marker genes for assigned T-cell populations (corresponding to Figure 6c and **6d**) grouped by CD4+, CD8+ and cycling cells. Marker genes are grouped by functional pathways (Lineage: CD4/CD8, Naive, Central Memory, Effector Memory, Actviation, Exhaustion, Cytotoxic, Treg, and Proliferation). **b.** Normalized CAR expression detection and grouped by expression decile bins. **c.** Pathway activity z-scores for exhaustion (*PDCD1*, *HAVCR2*, *LAG3*, *TIGIT*, *TOX*, *CTLA4*), activation (*CD69*, *CD38*, *ICOS*, *TNFSF4*, *TNFRSF18*), cytokine (*IL-2*, *TNF*, *IFNG*), memory (*TCF7*, *LEF1*, *CCR7*, *SELL*, *IL7R*), and TCR signaling (*CD3D*, *CD3E*, *CD3G*, *LCK*, *ZAP70*, *LAT*) markers across CAR expression deciles. Significance is determined by a linear mixed-effects model comparing mean gene set expression between cells in the top and bottom CAR-expression deciles with donor as a random effect (asterisk denotes p-value < 0.01). **d.** Alluvial plot of T cell subset composition changes across repetitive stimulation (RS) and IL-2 treatment conditions. Bar and ribbon widths reflect changes in population representation across each condition.

**Extended Data Fig 7.**
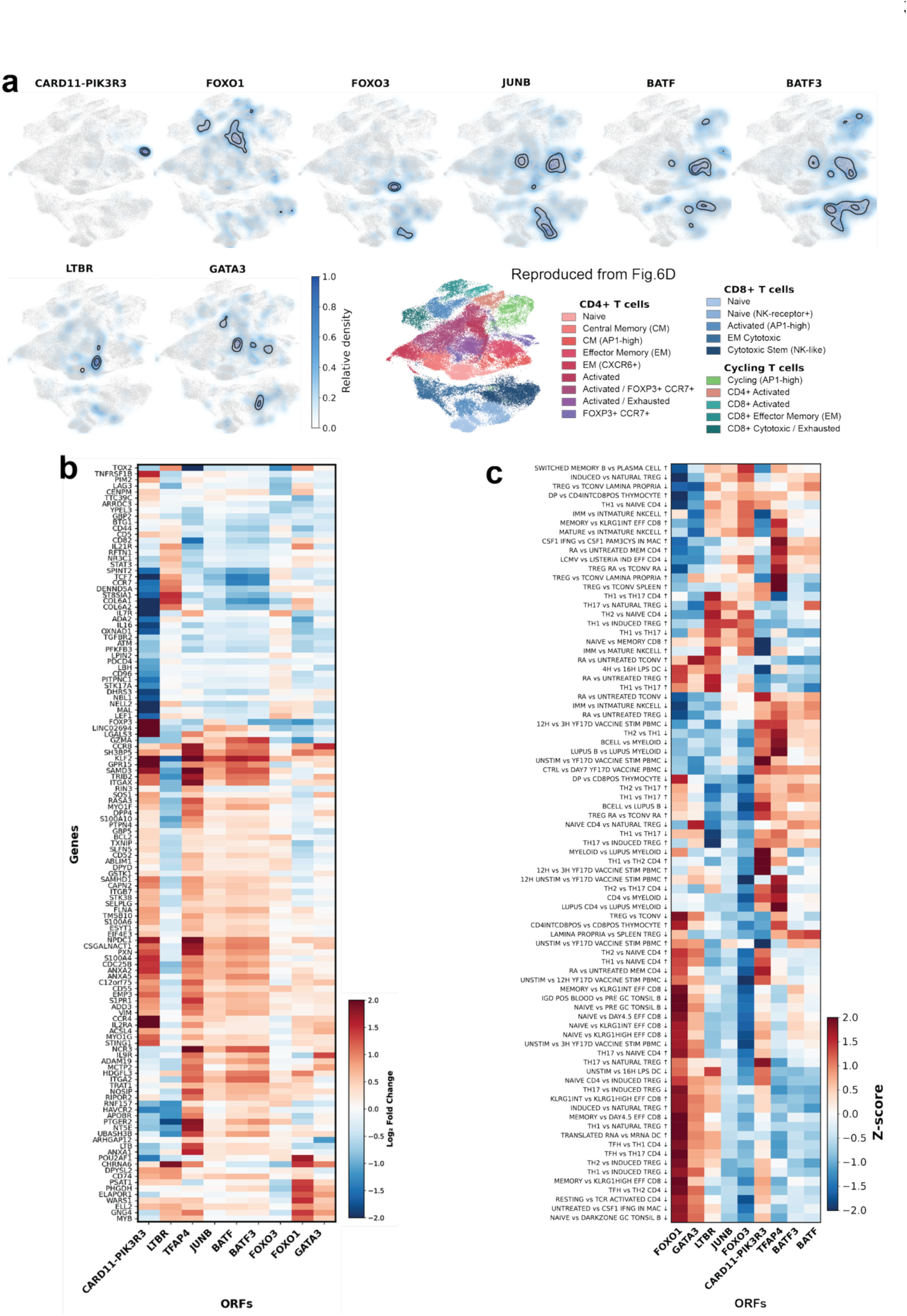
CoMPoSE measurement of ORF perturbation phenotypes identifies distinct. **a.** UMAP density plots showing spatial distribution of cells expressing individual ORFs. Light gray points represent all cells with a density gradient of ORF-expressing cells (white = low, dark blue = high). Black contour lines delineate regions of high enrichment (50% and 75% of maximum density). Figure 6d replotted for comparison. **b.** Clustered heatmap of differentially expressed genes in each ORF compared to mCherry control assigned cells. Shown as log_2_ fold-change for the top 125 most perturbed genes. **c.** Clustered heatmap of pathway enrichment for Immunesigdb pathways. Shown as z-score normalized pathway scores for each ORF.

